# “From Nutritional Patterns to Behavior: High-Fat Diet Influences on Inhibitory Control, Brain Gene Expression and Metabolomics in Rats”

**DOI:** 10.1101/2024.01.26.577449

**Authors:** Diego Ruiz-Sobremazas, Ana Cristina Abreu, Ángeles Prados-Pardo, Elena Martín-González, Ana Isabel Tristán, Ignacio Fernández, Margarita Moreno, Santiago Mora

## Abstract

**Background:** Impulsive and compulsive behaviors are associated with inhibitory control deficits. Diet plays a pivotal role in normal development, impacting both physiology and behavior. However, the specific effects of high-fat diets (HFD) on inhibitory control have not received adequate attention.

**Objectives:** This study aimed to explore how exposure to an HFD from postnatal day (PND) 33 to PND77 affects impulsive and compulsive behaviors.

**Methods:** The experiment involved 40 Wistar rats, half of which were male and subjected to the HFD or Chow diets. Several tasks were employed to assess behavior, including the Variable Delay to Signal (VDS), the Five Choice Serial Reaction Time Task (5-CSRTT), the Delay Discounting Task (DDT), and the Rodent Gambling Task (rGT). Genetic analyses were performed on the frontal cortex, and metabolomics and fatty acid profiles were examined using stool samples collected on PND298.

**Results:** The study revealed that rats exposed to the HFD exhibited heightened impulsivity in the VDS and in the 5-CSRTT, suggesting an increase in motor impulsivity. Notably, no significant effect was observed in the DDT. Surprisingly, the HFD-exposed group demonstrated reduced levels of impulsive-like behaviors, and a different decision making, in the rGT. Furthermore, abnormal gene regultation linked to brain plasticity and dopamine were identified in the frontal cortex. Metabolomics analysis of stool samples, collected in adulthood, indicated lower levels of fatty acids.

**Discussion:** These results suggest that HFD exposure during adolescence may create a lasting vulnerability to inhibitory control deficits, specifically in terms of motor impulsivity, and in gene expression as well as in metabolomics profile.

## 1. Introduction

Inhibitory control is defined as the ability to inhibit or control impulsive and/or compulsive responses (Bari & Robbins, 2013). Impulsivity and compulsivity are cardinal clinical features of several neuropsychiatric disorders like Attention Deficit Hyperactive Disorder (ADHD), Obsessive Compulsive Disorder (OCD), autism, or schizophrenia (American Psychiatric Association, 2013). In particular, impulsivity can be understood as a multifaceted phenomenon divided into different components: “impulsive action” as a failure in motor inhibition, “impulsive choice” as a tendency to accept small, immediate rewards over larger, delayed rewards, and “reflection impulsivity”, as an sensory evidence that has been insufficiently sampled (Dalley, Everit & Robbins, 2011; Evenden, 1999). Inhibitory control deficit is prominent during adolescence (Reichelt & Rank, 2017), and some factors such as diet (Vinuessa et al., 2016), substance abuse (Winters & Awwia, 2011) or even sport (Halson & Juliff, 2017) have been proposed to affect its normal development. However, the effects of some specific diets, like high fat diet (HFD), high sugar diet (HSD) or cafeteria diet (Caf), on the multifaceted nature of inhibitory control have not been fully studied.

HFD and HSD have been linked to impulsive behaviors in clinical studies (Lumley et al., 2016). Impulsive trait is connected with lower quality diet (Bénard et al., 2019). These results are more prone to appear in adolescence, which is a critical developmental period (de Andrade et al., 2016). In adolescence, impulsivity and diet variables are closely related (Smith et al., 2021; Lumley et al., 2016). Most studies aim to clarify the effects of obesity over different behaviors; however, few studies analyzed the effects of dietary manipulation over inhibitory control. Adams et al. (2015) exposed rats to two HFD and HSD. Increased motor impulsivity was found in subjects exposed to HFD assessed with the Five Choice Serial Reaction Time Task (5-CSRTT) compared with chow diet, while no effect in overall performance was found. Other studies analyzed choice impulsivity: no effect of HFD with 2:1 and 4:1 ratio of reinforcement was found (Garman, Setlow & Orsini, 2021); also, in HSD, no effect was reported (Wong, Dogra & Reichelt, 2017). Nevertheless, Robertson & Rasmussen (2017), that created an obesogenic phenotype, found an increased preference over Larger and Later (LL) rewards when subjects were exposed to a Caf diet. No studies have analyzed the effects of HFD/HSD or Caf consumption on impulsive decision-making. Dietary effects can also be detected even without creating an obesogenic phenotype (Leigh & Morris, 2020). For example, some differences were seen in the microbiome when a HFD is maintained for two weeks. In addition, long-term effects in plasmatic IL-1β were detected after 5 months since the HFD was exposed (Vinuesa et al., 2016). Additionally, some studies found a relationship between neuroinflammation and cognitive alterations (Leigh & Morris, 2020; Teixeira et al., 2017; Deshpande et al., 2019).

Furthermore, the activation of the immune system can also be secondary to diet (Cussotto et al., 2018). Some studies found an increased in pro-inflammatory molecules when HFD (Vinuesa et al., 2016), HSD (Beilharz, Maniam & Morris, 2016) and Caf (Cussotto et al., 2018) is presented. Additionally, Zeeni et al. (2013) found that when rats were exposed to a chronic variable stress, no difference was found in the high palatable diet groups; this diet was associated with a reduction response to chronic stress. In addition, Shin et al. (2019) found an increase IL-1β in serum after generating an obesogenic phenotype. However, the effects of these diets are not focused in the gut-microbiome only; they can also affect the mesolimbic and fronto-striatal pathways via activating DA neurons (Altherr et al., 2021). Besides, some proteins such as Brain-derived neurotrophic factor (BDNF) were reduced in male rats with four week HFD while remaining intact in females (Liu et al., 2014). The existent relationship between BDNF and diet is still unknown. Long and acute exposure (2-8 months) to HFD modifies BDNF concentrations, while short exposure (20-42 days) does not seem to affect concentrations (Fadó et al., 2022).

Brainstem and striatum seem to be two biological targets for high-fat diets effects (Alsiö et al., 2014). Specifically, in the striatum we can find dopamine (DA) receptors 1 and 2 (DRD1, DRD2) which cohabitate with cannabinoid receptor 1 (CB1), which is closely related to food regulation and eating disorders like obesity or binge-like eating (Bourdy et al., 2021; Volkow, Wise & Baler, 2017; Scherma et al., 2014). Regarding DA, up-regulation of the DRD1 in the amygdala was found in rats exposed to HFD throughout adolescence (Fülling et al., 2020); in brainstem, an increase in relative expression in DRD1 and a reduced relative expression of DRD2 was detected. However, other authors (Leeuw van Weenen et al., 2009) found no differences in genes related to the DA system (*DRD1*, *DRD2*, *TH* and *DAT*). In addition, no differences were present in *D1* and *D2* but in *COMT* in rats exposed to HFD; this difference was located in obesity-prone rats (Bourdy et al., 2021). One part of the mesolimbic circuit is the Nucleus Accumbens (Nacb). The Nacb shell motivates consumption of dietary fat in rats, and when D1Rs are inhibited in the lateral shell, a reduction in fat consumption can be perceived (Joshi et al., 2021). The endocannabinoid system is formed by CB1 and CB2 receptors. Rojo et al. (2013) found an increase in CB1 stimulation in the prefrontal cortex after 4-12 HDF consumption without an increase in receptor density, but no differences were present in chronic diet consumption (16-20 weeks). Regarding the Nacb, no differences were seen in *CB1* or *CB2* expression (Bourdy et al., 2021). In the case of BDNF, sexual dimorphism was noticed in the hippocampus after 28 weeks under HFD (Prochnik et al., 2022). BDNF levels are able to change even with a 24-hour HFD consumption, finding differences between sexes. When exposure time rises, those differences change until differences between sexes are more pronounced (Liu et al., 2014). However, fat diets can also affect glutamatergic transmission in the hippocampus, specifically down-regulating NMDA receptors like *Grin2a* and *Grin2b* (Fernández-Felipe et al., 2021). There is some overlap between the biological targets of HFD and the brain circuit related to inhibitory control. The fronto-striatal pathway is related with inhibitory control (Morein-Zamir & Robbins, 2015) and it is formed by the prefrontal cortex (PFC), orbitofrontal cortex (OFC), anterior cingulate cortex (Cg1), infralimbic, the Nacb core (NacbC) and shell (NacbS), and some nuclei of the ventral tegmental area (VTA) (Dalley, Mar, Economidou & Robbins, 2008).

Thus, the present study aims to investigate the putative relationship between HFD consumption at a young age, and inhibitory control deficits in adulthood, using premature responses as a measure of impulsivity while perseverative responses were used as a measure of compulsivity, as well as determining which neurochemical changes may be related with those deficits. Different paradigms, such as the Variable Delay to Signal (VDS), 5-CSRTT, DDT and rodent Gambling Task (rGT), were used to screen the three components of inhibitory control. Genetical analyses were performed using RT-qPCR in frontal cortex, while metabolomics analyses were performed in stool samples using ^1^H NMR and GC-FID analyses; both collected at the last stage of the protocol. Our hypothesis is that rats exposed to HFD during a critical developmental period will show impaired inhibitory control-related measures compared with the chow-fed group. Furthermore, we expect differences in some genes related to neurotransmission systems in the frontal cortex. Also, we expect differences in the metabolomics profiles and in the fatty acids of our groups.

## 2. Material and methods

### 2.1. Subjects

40 Wistar rats (20 male and 20 female; ENVIGO, Barcelona, Spain) were used in the present study. They arrived at the lab on the postnatal day 21 (PND21) and were housed in groups of four rats per cage (57 x 35 x 20 cm), at 22 ± 1 °C and under a 12:12-h inverted light-dark cycle with lights off at 09:00. Environmental enrichment (PVC and wooden blocks) was added to their home cages. Food and water were provided *ad libitum*. At arrival, eleven days of habituation to the environment took place, after which handling was performed daily along with body weight gain, food and water intake assessment (described in baseline consumption assessment). After this, body weight control was performed once per week. Assignment to each of the experimental groups (Chow or Cheese) was done randomly. Once the diet manipulation was over (PND77), rats were fed *ad libitum* until PND96. Food deprivation started with the objective of achieving an 85% of their weight at PND96 until sacrifice PND298.

All the procedures were conducted in agreement with Spanish Royal Decree 55/2013 on the protection of experimental animals and European Directive (2010/63/EU), and were approved by the University of Almería Research Committee. All the researchers show commitment to the three Rs principle.

### 2.2. Experimental design

Subjects were assigned to one of two experimental conditions: chow (*n* = 20) or Cheese (*n* = 20). After eleven days of habituation to the laboratory (PND21-PND32), a test of basal consumption (PND33) was performed, after which exposure to HFD began (PND33-PND77). On the initial days of dietary manipulation, a test of HFD-related consumption was performed (PND34-37). Once in adulthood, dietary manipulation ended and, after two weeks of stabilization and chow-based diet, food restriction was gradually performed until animals reached 85% of their previous bodyweight. From PND96, a normocaloric diet was used for maintaining subjects at 85%. Behavioral assessment started at PND112 and finish at PND297 (Figure 1). Brain dissection and stool collection were performed at PND298.

**Figure 1.**
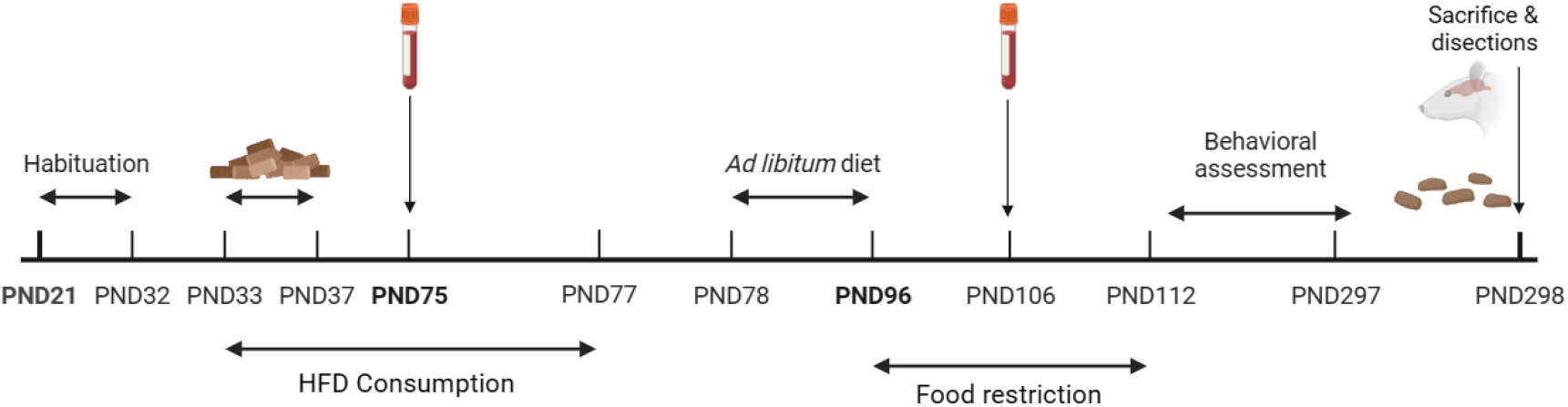
Experimental and behavioral procedure illustrated in a timetable. After arriving at postnatal day 21 (PND21) and a period of habituation to the laboratory (PND21-32), a basal chow test consumption between PND33 and PND37 was conducted, and between PND33-77 the High-Fat Diet (HFD) was given to the rats. Within the HFD time, a blood collection test was done at PND75. After the HFD was complete, *ad libitum* diet was given from PND78 until PND96. From PND96 until PND112 diet restriction until rats achieve 85% of their weight. Another blood collection was done at PND106. The behavioral assessment followed, starting at PND112 until PND297. At the end, rats were sacrificed at PND298 where brains and stool samples were collected.

### 2.3. High-Fat Diet (HFD) protocol

In accordance with previous literature (Lalanza & Snoeren, 2021), highly caloric, high-fat diet was provided based on commercial cheesecake (Postres Reina, Caravaca de la Cruz, Spain), with the following energetic and macronutrient specifications (*per* 100 g): 800 kJ/191 kcal as energetic value (10%); 9.9 g fats (14%) of which 6.1 saturated (31%); 21.4 g carbohydrates (8%) of which 18.5 g (21%) sugar; 4.1 g of proteins (8%); 0.2 g of salt (2%). Daily quantities were chosen based on the recommendations by Leigh et al. (2019), with a total of 1kJ per g for each rat, and adjusted based on bodyweight evolution. Laboratory diet remained *ad libitum* for both groups during dietary manipulation.

### 2.4. Behavioral assessment

Task organization was set in order not to interfere with each other. To prevent this from happening, behavioral assessment started with a light cue task (Variable Delay to Signal; VDS) followed by a lever cue task (Delay Discounting Task; DDT); both tasks are different in response (light response in VDS and lever-press response in DDT). After these tasks, the 5 choice serial reaction time task (5-CSRT), a light response task, was conducted; afterwards, the rodents Gambling Task (rGT) was conducted. A minimum of one-week washout was left between tasks. All the behavioral assessment was performed during the dark/active phase.

#### 2.4.1. Apparatus

The behavioral tests were performed in one set of six and other of eight operant-conditioning chambers (MED Associates) measuring 32 cm long x 25 cm wide x 34 cm high, with stainless-steel grille floors. Neither the modules/wall panels used for VDS, DDT and 5-CSRT (set 1, six chambers) were changed between tasks. A 5-CSRT task panel wall, consisting of five contiguous square holes (2.5 cm), a height of 2 cm above the grilled floor and 2.2 cm deep, was used for VDS and 5-CSRT; a detailed description can be found in Moreno et al. (2010, 2012). For DDT, two retractable levers were available in the opposite wall of the operant chambers; details are available in Cardona et al (2006, 2011). In order to minimize any possible interference between tasks, the order was as follows, VDS (nosepoke) - DDT (lever press) - 5-CSRTT (nosepoke) - rGT (nosepoke). Each task commenced one week after the previous one. The scheduling and recording of experimental events were done by a Med PC computer and a specific commercial software (Cibertec SA, Spain). Specific details of each task can be found in the Supplemental Material, Behavior analyses.

### 2.5. Biochemical Analyses (RT-qPCR & ELISAs)

Blood samples were collected in two different moments. First, while in HFD at PND75; second, before behavioral analyses (PND106). After completion of all tasks, all animals were deeply anesthetized with isoflurane and sacrificed with fast decapitation. Prefrontal cortex and nucleous accumbens were quickly dissected and stored separately into RNAse-free tubes (1.5 mL). Also, stool samples were recollected from each rat. All samples were immediately fresh-frozen to avoid RNA degradation. All materials required were autoclaved and cleaned with RNAse ZAP (Invitrogen). Samples were stored at −80°C until use. RTqPCR was performed in the prefrontal cortex and ELISA’s analyses were conducted with serum. The specific procedures of each biochemical assessment can be found in the Supplemental Material, Biochemical analyses.

### 2.6. ^1^H NMR Analyses

The detailed protocol that was used for the ^1^H NMR analyses can be found in the Supplemental Material, Metabolomic analyses.

### 2.7 GC-FID analysis

The fatty acid profile and content in 10 mg of each fecal sample were determined by gas chromatography (Agilent Technologies 6890 N Series Gas Chromatograph, Santa Clara, CA, USA) after direct transesterification (Rodríguez-Ruiz et al., 1998).

### 2.7. Statistical analyses

Baseline weight gain was analyzed by a one-way analysis of covariance (ANCOVA), with group (Chow vs Cheese) as between-groups factor and sex (male vs female) as covariable. Bodyweight evolution during consumption test was analyzed by a two-way repeated measures (RM) ANCOVA, with group as between-groups factor, day (1-5) as within-subjects factor, and sex as covariable. Baseline water and Chow consumption were analyzed by a one-way ANCOVA, with group as between-groups factor and sex as covariable. Consumption test was analyzed by a two-way RM- ANCOVA, with group as between-groups factor, day as within-subjects factor, and sex as covariable. Behavioral assessment was analyzed by a two-way RM-ANCOVA (for the learning) and one-way ANCOVA (for the test) in the VDS with group and session as a between- and within-subjects factor respectively. In the DDT and 5-CSRTT, two-way RM-ANCOVA was conducted with group as a between factors and delays (DDT) or stimulus durations (5-CSRTT) as a within-factor. In the rGT, one-way RM-ANCOVA was conducted with group as a between factor. RT-qPCR results were analyzed with t-test between groups in both sexes with group as a between-factor. ELISAs results were analyzed via two-way RM-ANCOVA with group as a between-subjects and time assessment as a within-subjects factor. In all ANCOVAs, sex was set as the covariable. If this covariable reached significant levels, a split analysis was conducted in order to fully understand the data dynamics. Post hoc analyses were performed, when necessary, with Bonferroni corrections. Outlier values were calculated with GraphPad Prism tool and removed if present. Statistical significance was set at p < 0.05, and effect size is reported when appropriate: Partial eta-squared values are reported and considered as small (0.01), medium (0.06), or large (0.14) following Cohen (1988) recommendations. All analyses were carried out using Statistica^®^ software (Statsoft, version 6.0) and JASP^©^ (University of Amsterdam, version 0.14.1). Graphs were created using GraphPad Prism (San Diego, California, USA) v8.0.0, while images were designed using Biorender.

For NMR data, multivariate data analysis was performed on the obtained dataset using SIMCA-P software (v. 17.0, Umetrics). Exploratory and unsupervised analysis as principal component analysis (PCA), and supervised models as partial-least squares discriminant analysis (PLS-DA) were applied by scaling data to Unit Variance. Scores and loadings plots were generated for both models. PLS-DA models were validated by means of their goodness-of-fit (R^2^) and goodness-of-prediction (Q^2^) cumulative value, together with the CV-ANOVA parameter validation (at the level of significance of *p* < 0.05) to test the accuracy of the model. Loadings containing important metabolites for predictive models were evaluated by generating the variable importance in projection (VIP) plot and selecting those superior to 1. Fold-changes for discriminant metabolites among groups were estimated. Wilcoxon rank-sum tests were applied to determine the significance of the metabolites (*p* < 0.05) employing the online tool MetaboAnalyst. All the metabolites that significantly changed between groups regardless of the FC value were considered.

## 3. Results

### 3.1. Baseline weight, chow and water consumption

No differences were detected in body weight concerning conditions-to-be-assigned in the first day; however, the expectable sex effect (F_1, 37_= 47.299; *p* < 0.001; partial η^2^= 0.561) and an overall day effect (F_1,148_ = 256.891; p < 0.001; partial η^2^ = 0.874) were present. No differences between groups were shown, but a predicted sex effect was shown (F_1,37_ = 54.576; p < 0.001; partial η^2^ = 0.596). Day*group (F_1,148_ = 4.202; p < 0.001; partial η^2^ = 0.102) and day*sex (F_1,148_ = 8.978; p < 0.001; partial η^2^ = 0.195) interactions were observed. In baseline privation day, only a sex effect existed (F_1, 37_ = 304.860, p < 0.001, partial η^2^ = 0.892).

In the baseline chow consumption test, a group (F_1, 37_ = 6.823; p < 0.05; partial η^2^ = 0.156) and sex effect (F_1,37_ = 14.209; p < 0.01; partial η^2^ = 0.185) were detected. Concerning chow consumption through the assessments, a significant day (F_4, 148_ = 48.402; p < 0.001; partial η^2^ = 0.567), sex (F_1, 37_ = 78.204; p < 0.001; partial η^2^ = 0.679) and group effect (F_1, 37_ = 256.891; p < 0.001; partial η^2^ = 0.874) were present. The day*group (F_1,148_ = 26.682; p < 0.001; partial η^2^ = 0.419) and the day*sex (F_1,148_ = 10.604; p < 0.001; partial η^2^ = 0.223) interactions were observed. Concerning baseline water consumption, both group (F_1, 37_= 5.293; p < 0.05; partial η^2^ = 0.125) and sex (F_1, 37_= 7.474; p < 0.01; partial η^2^ = 0.168) effects were perceived. An overall day effect was found (F_4, 148_= 15.947; p < 0.001; partial η^2^ = 0.301), as well as group (F_1,37_ = 41.159; p < 0.001; partial η^2^ = 0.527) and sex (F_1,37_ = 20.200; p < 0.001; partial η^2^ = 0.353) effects regarding water consumption across all test days, but no interaction was obtained (Figure 2).

**Figure 2.**
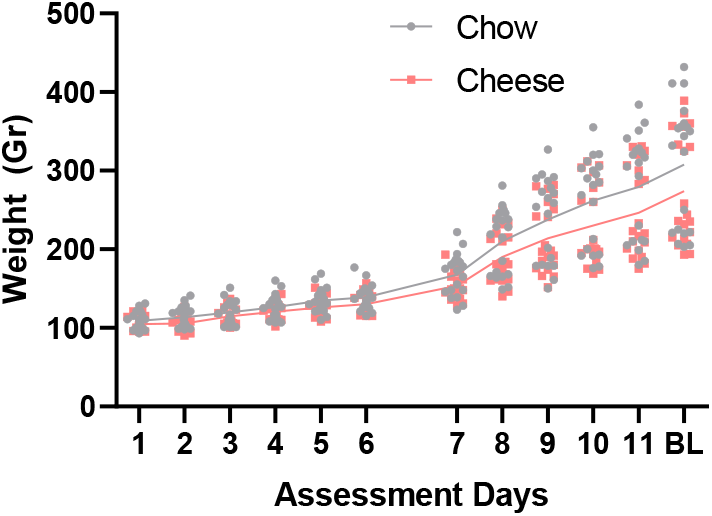
Weight evolution across all test days until Baseline (BL) privation day. The period shown corresponds to the period where the rats were fed with HFD. Comparisons were performed using Bonferroni correction. Individual data is represented and line connects mean group values. Total n =20 in each group. Numbers from 1 to 6 represent weight evolution daily, numbers between 7 and 11 represent body weight assessed every week.

### 3.2. Variable Delay to Signal

#### 3.2.1. VDS training performance and inhibitory control

All groups showed appropriate learning, reducing total session time (F_9,333_ = 16.976; p < 0.001; partial η^2^ = 0.315; post hoc comparisons revealed that S1-S3 were different between them and between the other sessions; p < 0.05 in all comparisons) and increasing total correct responses (F_9,333_ = 5.297; p < 0.001; partial η^2^ = 0.125; session one was different with the other sessions with p < 0.001 using Bonferroni correction), reducing mean (F_9,333_ = 14.256; p < 0.001; partial η^2^ = 0.278; post hoc revealed that session 1 and 2 were different from the others with p < 0.001) and total (F_9, 333_= 16.976; p < 0.001; partial η^2^ = 0.315; post hoc comparisons revealed that session 1 to 3 were different to the others with p < 0.001) latency response, and total (F_9,333_ = 10.864; p < 0.001; partial η^2^ = 0.227; same post hoc results were found. Sessions 1 and 2 were different from the others with p < 0.001) but not mean latency to reward. No effect of sex nor group was found (Figure 3). Rats also tended to do less omissions across sessions (F_9, 333_= 13.890; p < 0.001; partial η^2^ = 0.273; sessions 1 and 2 had more omissions compared to the others with p < 0.001). A session*sex interaction was found (F_9, 333_ = 2.387; p < 0.05; partial η^2^= 0.061) in total session time, however, no effect of sex was evidenced.

**Figure 3.**
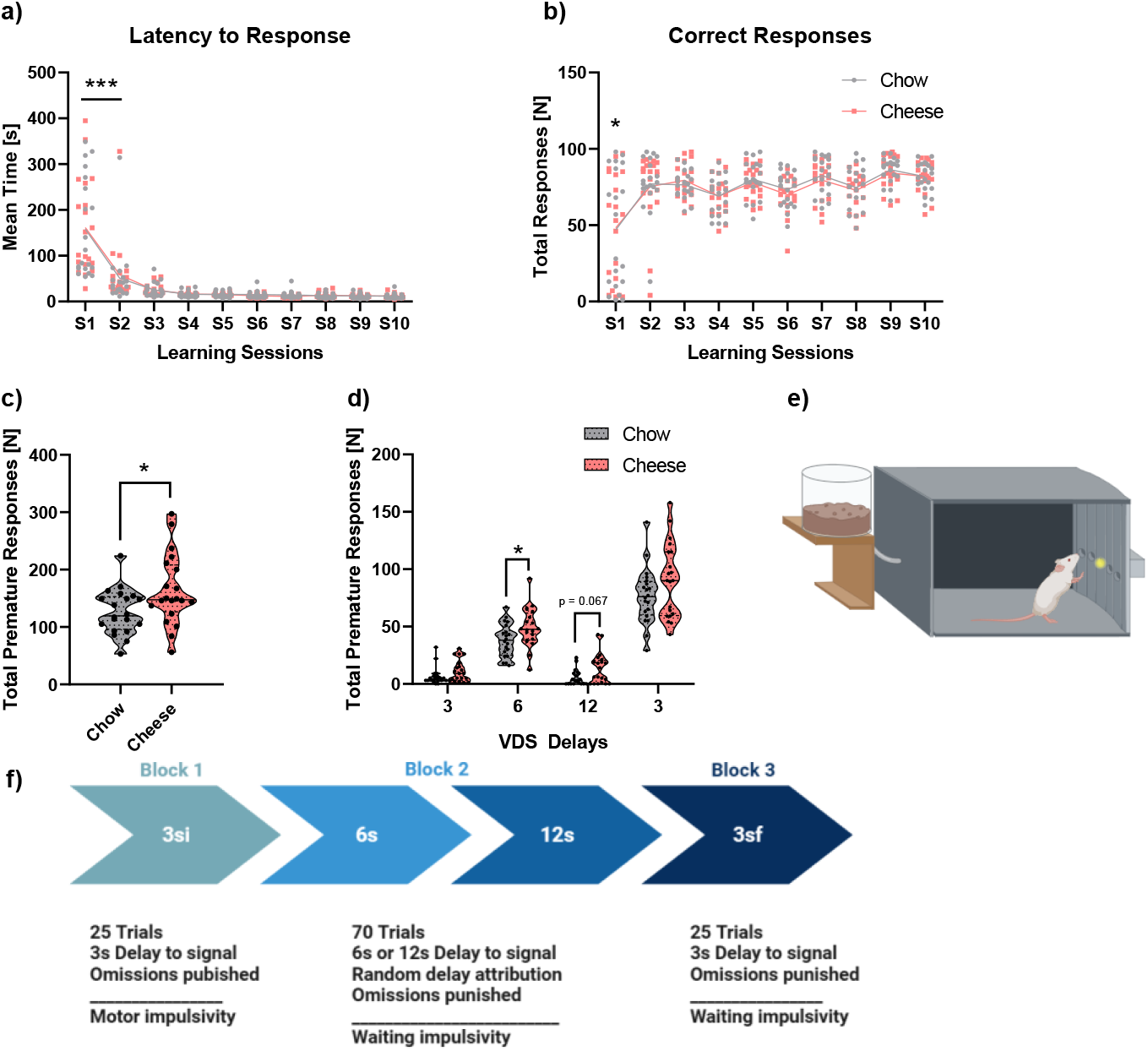
VDS training and test performance representation. In the upper part, mean latency to response across all training sessions (a) is depicted while in b) correct response through learning sessions are shown. In the lower part, total premature responses (c) and total premature responses per minute in the test session are shown. In part e) a graphical example of the operant box used. Lastly, in part f) a graphical resumee of the task is shown. Total n in each group was 20. Data is represented as individual values with lines connecting means. In the lower part, violin graphs are depicted, with mean and quartiles; * p < 0.05; *** p < 0.001. Differences in graphs a) and b) correspond to session differences, not diet nor sex.

No effect was found in premature responses: although an overall session effect (F_9,333_ = 8.579; p < 0.001; partial η^2^ = 0.188), no group or sex effects were detected. Moreover, in prematurity response rate an overall session effect was observed (F_9,333_ = 5.651; p < 0.001; partial η^2^ = 0.132), no group or sex effects were found. Rats did more premature responses throughout all training sessions. Similarly, no effect existed in perseverative responses: although a session effect was found (F_9,333_ = 4.223; p < 0.001; partial η^2^ = 0.102), no group nor sex effects were detected. Rats made more perseverative responses in the first three training sessions.

#### 3.2.2. VDS Test performance and inhibitory control

No effect was found in correct responses, omissions, total response latency, mean response latency, total latency to reward or mean latency to reward. On the contrary, a strong effect of group was found in total premature responses (F_1,35_ = 4.872; p < 0.05; partial η^2^ = 0.122) (Figure 3). When analyzing the different delays, an effect of group was detected in the 6 seconds delay (F_1,35_ = 4.198; p < 0.05; partial η^2^ = 0.107), as well as a trend close to significance in the 12 second delay (F_1,35_ = 3.885; p = 0.057; partial η^2^ = 0.1), but no effect were noted in the others. No sex effect was perceived in total premature responses in any of the VDS blocks. Rats with HFD consumption tended to do more premature responses regarding total premature responses and premature responses in 6- and 12-second delay.

### 3.3. Five Choice Serial Reaction Time Task (5-CSRTT) training performance and inhibitory control

All rats learnt the task correctly. No differences were found according to the sessions required to reach each criterion. However, a significant difference was observed in the total sessions required to achieve SD1 criteria (F_1, 29_= 4.989, p < 0.05, partial η^2^ = 0.147). Control group needed more sessions to achieve criteria. The ANCOVA revealed no differences in any learning variables in the three consecutive sessions required to achieve SD1 criteria. However, the ANCOVA revealed that the Cheese group did more premature responses than control (F_1, 31_ = 4.637; p < 0.05; partial η^2^ = 0.128). No effect was found for perseverative responses (Figure 4).

**Figure 4.**
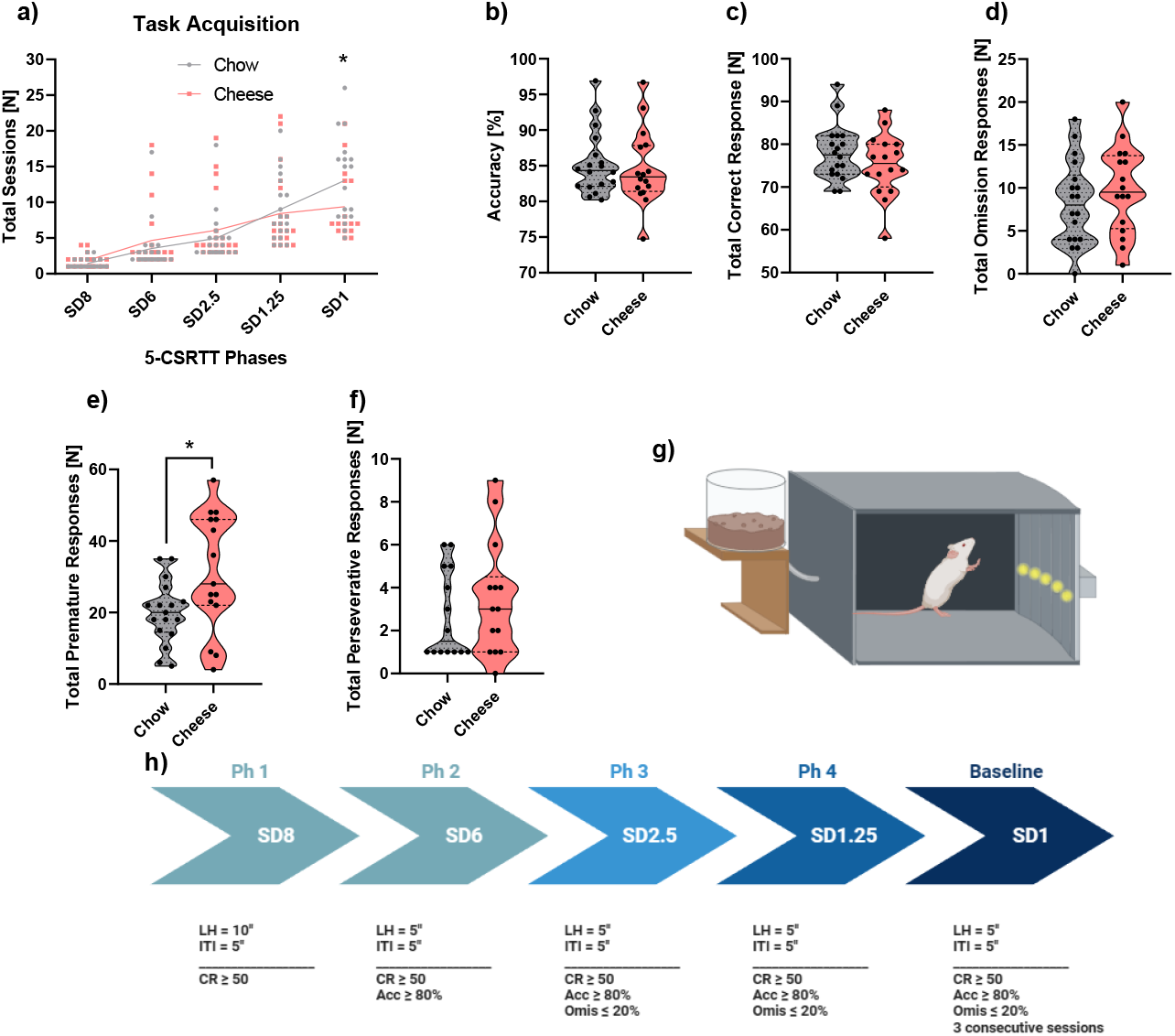
Graphical image of the 5-CSRTT learning and baseline inhibitory control measures. In part a) total sessions to achieve baseline condition (SD1) is shown; line connects mean in every session. In part b), c) and d) task learning is shown as accuracy, correct responses and total omission responses respectively. In part e) total premature responses are depicted and in part f) total perseverative responses is shown; these graphs are violin plot where mean and quartiles are represented. In part g) a graphical example of the operant box used. Lastly, in part h) a graphical resume of the task is shown. Individual data is represented as well as mean ± SEM. * p < 0.05. Sample sizes were as follows: n = 16 for Cheese and n = 18 for Chow. * refer to group differences (Chow vs Cheese).

### 3.4. **DDT**

The RM-ANCOVA revealed a large delay effect across all delays (F_4, 128_ = 27.691; p < 0.001; partial η^2^ = 0.461). No effect of group was found, nor sex effect in any comparison. Rats showed more mean choices for the LL during the DDT (Figure S1). All the animals chose the LL when the delay is 0 or close (5 seconds), but when this increases, they tend to choose the SS.

### 3.5. **rGT**

All rats learnt the task accordingly. A significant effect of group was found in percent premature responses (F_1, 35_ = 8.381; p < 0.01; partial η^2^= 0.235), sex also reached significant levels (F_1, 35_ = 7.462; p < 0.01; partial η^2^ = 0.176), but its interaction was not significant. The Cheese group did less premature responses than chow, and males did higher compared with females (Figure 5).

**Figure 5.**
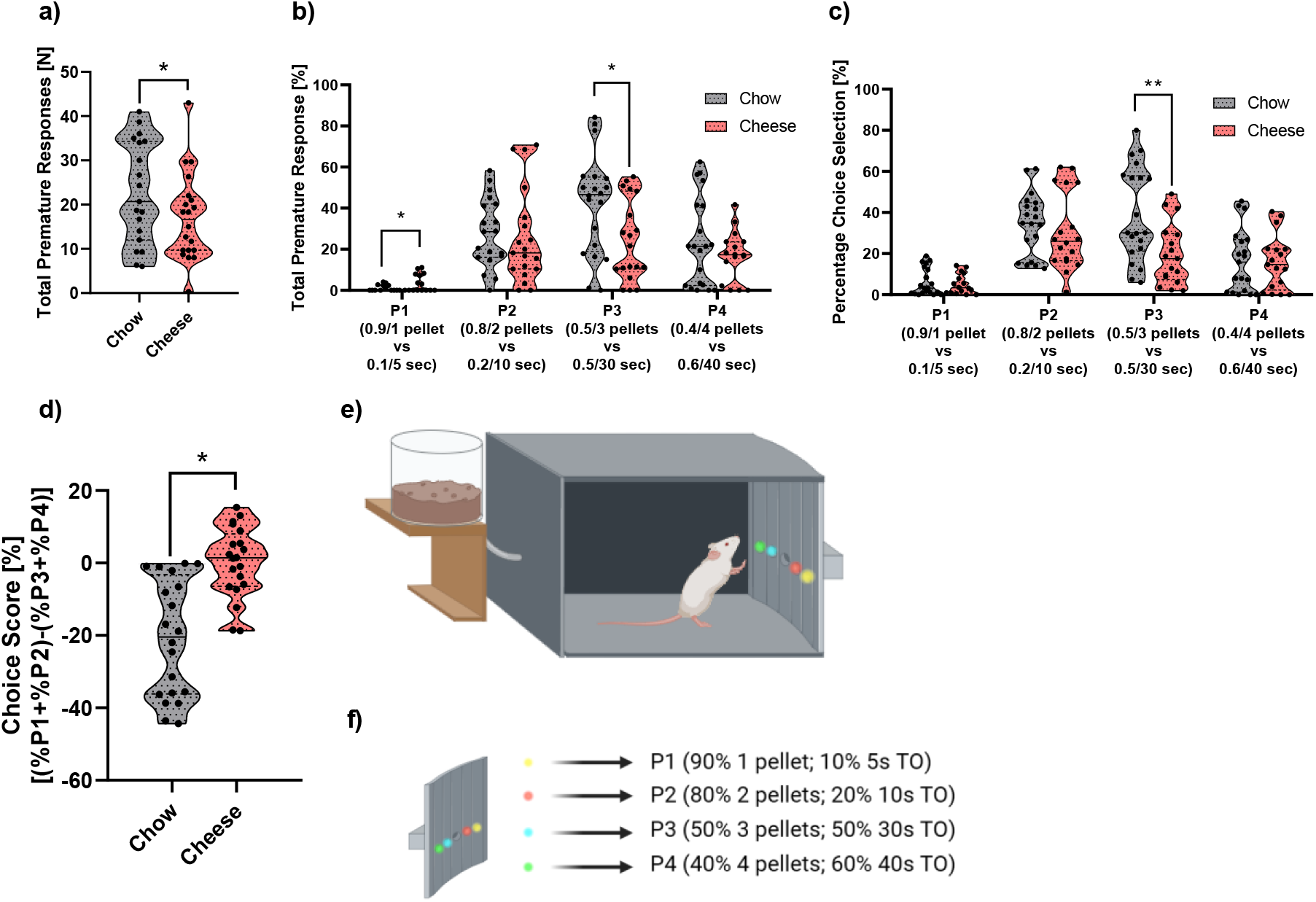
Graphical representation of rGT’s variables. In the upper part, total premature reponse (a) and total percentage premature response (b) is depicted. In addition, Percentage choice selection (c) and percentage choice score (d) are shown. Sample size is as follows: in graph a) (19 for chow and 20 for Cheese), in part b) (P1 (n chow = 14; n Cheese = 15), P2 (n chow = 19; n Cheese = 19), P3 (n chow = 20; n Cheese = 19) and P4 (n chow = 20; n Cheese = 15). In graph c) (P1 (n chow = 20; n Cheese = 16), P2 (n chow = 18; n Cheese = 18), P3 (n chow = 20; n Cheese = 17), P4 (n chow = 19; n Cheese = 17), in part d) is 20 in each group. Individual plots were shown when possible, mean ± SEM is depicted in every image. * p < 0.05, ** p < 0.01

The premature responses were also analyzed for every possible response. We found a trend to significance in P1 (F_1, 26_ = 3.597; p = 0.069; partial η^2^ = 0.122) without any sex effect and a group significance in P3 (F_1, 35_ = 4.709; p < 0.05; partial η^2^ = 0.119), being the Cheese the less impulsive, with a trend to significance in the sex (F_1, 35_ = 3.879; p = 0.057; partial η^2^ = 0.100; Figure 5). Cheese group showing less percent premature responses (M = 25.272; SD = 19.233) than chow group (M = 40.546; SD = 23.058) and females (M = 25.978; SD = 18.569) higher than males (M = 39.839; SD = 23.722). Same results in P3 were found in percent choice selection for group (F_1, 34_ = 7.357; p < 0.01; partial η^2^ = 0.178). Sex did not reach significant levels. No effect was detected for perseverative responses in total or in percent.

In addition, a significant difference was evidenced in percent reinforced trials (F_1, 30_ = 6.365; p < 0.05; partial η^2^ = 0.175), where Cheese got more reinforced trials than control (Cheese; M = 64.214; SD = 7.113; Chow; M = 58.279; SD = 3.232). A trend for group was observed in total latency (F_1, 32_ = 4.018; p = 0.054; partial η^2^ = 0.112), being the Cheese the group with higher latencies (Cheese; M = 59412.627; SD = 27095.943; Chow; M = 44389.019; SD = 12350.852). No difference was detected for punished trials.

To end up with, we detected a strong effect of group on percent choice index (F_1, 37_ = 5.523; p < 0.05; partial η^2^= 0.130). Cheese group had higher index than Chow.

### 3.6. RT-qPCR Gene Expression

Independent t-tests were performed according to sex to be able to use the chow groups (chow-male and chow-female) as control. In the case of male rats, differences were found in *BDNF* Fold Change (p < 0.01; d = −2.593), in *CB1* (p < 0.05; d = −1.613) and in *DRD1* (p < 0.05; d = −2.018). No differences were detected in *DRD2*, *GAD1*, *TNF-*α and *TYRO* (Figure 6) fold change in PFC. Regarding female rats, differences were obtained in *BDNF* Fold Change (p < 0.05; d = 2.620), but no differences were noticed in *CB1*, *DRD1*, *DRD2*, *GAD1*, *TNF-*α or *TYRO* (Figure S2).

**Figure 6.**
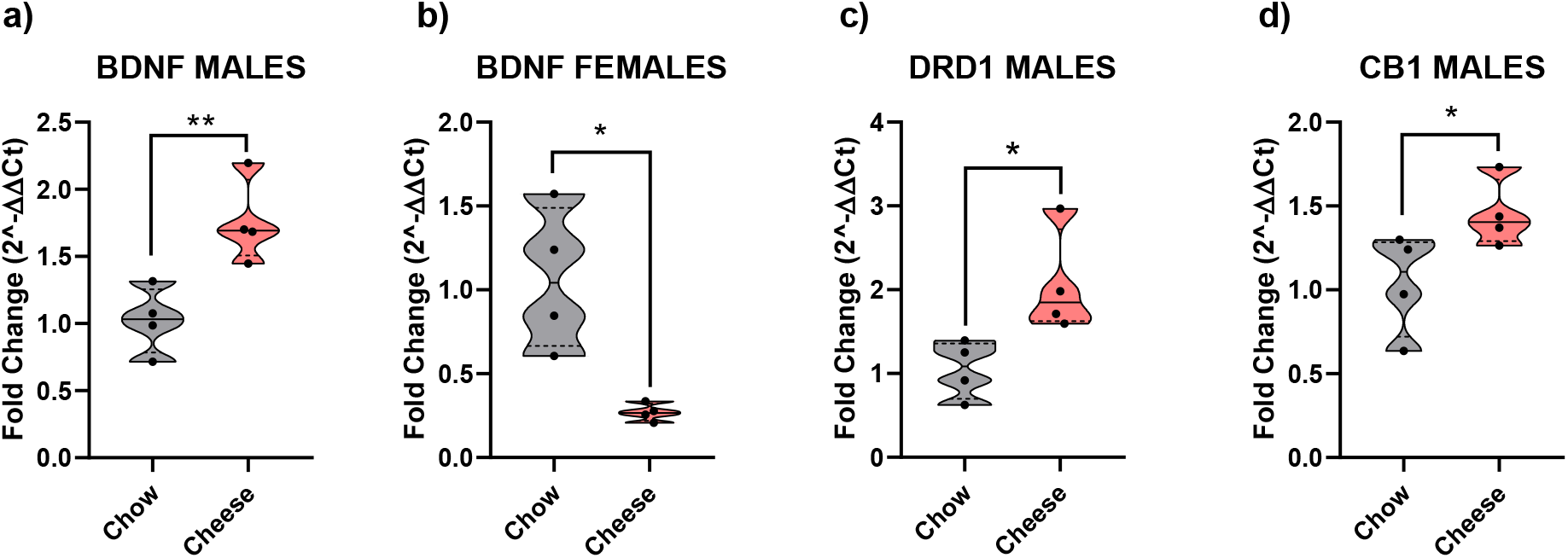
Visual representation of the genes that achieved significative differences in the t-students. * p < 0.05; ** p < 0.01. Total n = 4 in each group

### 3.7. CORT, leptin and TNF-α serum level analysis

Regarding leptin, the RM-ANCOVA revealed differences between time measures (F_1, 22_ = 19.030; p < 0.001; partial η^2^ = 0.464). The covariable also reached significance levels (F_1, 22_ = 17.095, p < 0.001: partial η^2^ = 0.437); when sex was included as an intra-subject factor, differences were also detected (F_1, 21_ = 13.734; p < 0.001; partial η^2^ = 0.395). Males and females were different in leptin concentration in the first assessment (F_1, 24_ = 11.587; p < 0.01; partial η^2^ = 0.345) and in the second assessment (F_1, 24_ = 18.975; p < 0.001; partial η^2^ = 0.463) (Figure 7).

**Figure 7.**
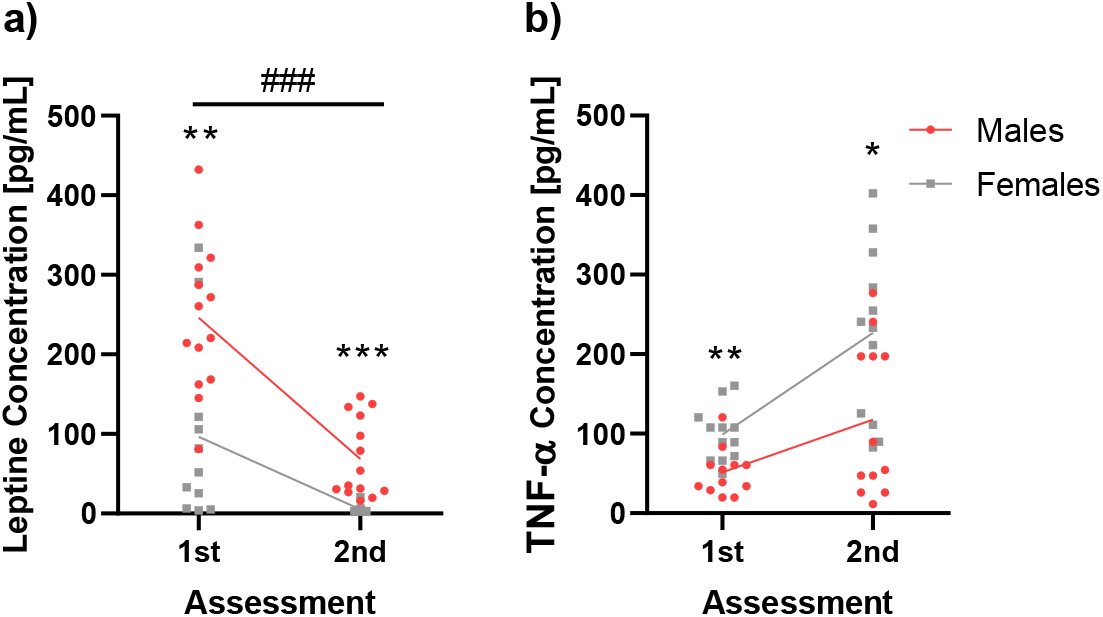
Graphical representation of leptine (a) and TNF-α (b) in serum between the two assessments. Individual values are shown in every assessment and in every condition. Sample size is as follows: n = 14 for males and n = 13 for females in graph a) and n = 12 for graph b). * p < 0.05, ** p < 0.01, *** p < 0.001, ### p < 0.001. * represents group differences, while # represents assessment differences.

TNF-α levels were equal between assessments and groups. In this RM- ANCOVA, the covariance reached significance levels (F_1, 22_ = 13.309; p < 0.001; partial η^2^ = 0.377). When sex is included as an intra-subject factor, sex also reached significant differences in the first assessment (F_1, 24_ = 11.303; p < 0.01; partial η^2^ = 0.350) and in the second assessment (F_1, 24_ = 4.625; p < 0.05; partial η^2^ = 0.180) (Figure 7).

### 3.8. NMR and GC-FID metabolic profiles

On the one hand, the NMR analyses showed several essential and non-essential amino acids (valine, leucine, isoleucine, alanine, threonine, glutamate, aspartate and glycine). Furthermore, organic acids (succinate, gumarate, formate, choline, betaine, glucose, bile acids, short-chain fatty acids [SCFA; acetate, propionate and butyrate]), that serve as an indicator of microbial and metabolomic processes, were detected. All NMR detected metabolites are explained in the Table S1.

On the other hand, the GC-FID analyses revealed 13 fatty acids (FA), from which 7 were saturated (palmitic, capric, and lauric acid the major ones), 3 monounsaturated (aicosenoic acid the most frequent, followed by oleic and vaccenic acids), and 3 polyunsaturated fatty-acid (PUFA) chains from linoleic acid. Specifically, saturated FA and PUFAS were found in a higher ration in female stool samples (Figure 8; Table S2). Furthermore, other acids (cis-9,12-hexadecatrienoic and cis-6,9,12-hexadecatrienoic) were present in female stool samples while remained undetected in male stool samples.

**Figure 8.**
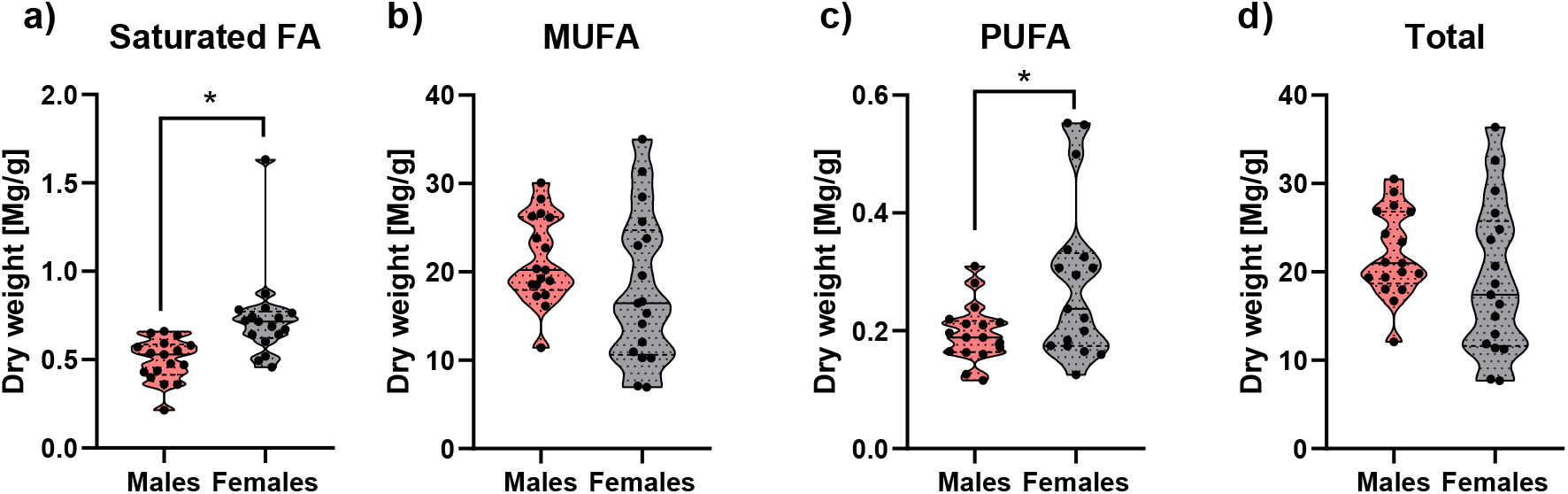
Boxplots of FA concentrations, distributed in saturated, monounsaturated, and polyunsaturated FA, and total FA, and detected by GC-FID in male and female feces. Differential concentrations determined by unpaired t-test (*p* < 0.05) between male and female feces were marked with *. Total n in each group is 17.

**Figure 9.**
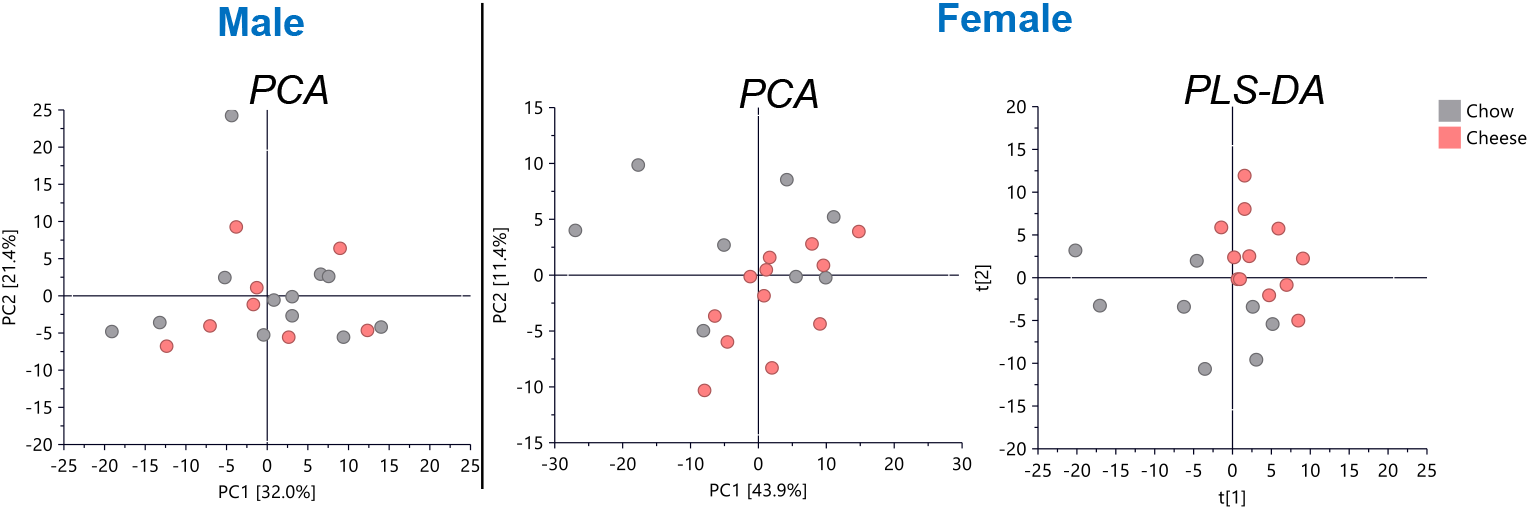
PC1/PC2 PCA models applied to ^1^H NMR data of fecal extracts from male and female rats. A valid PLS-DA model showing discrimination between Chow and Cheese groups was only found in female fecal extracts (R^2^X = 0.61, R^2^Y = 0.994, Q^2^ = 0.514, CV-ANOVA = 0.22). Total n in the Male PCA is as follows: Chow (n= 12), Cheese (n = 8); Female PCA n is as follows: Chow (n= 8), Cheese (n= 12); Female PLS-DA n is: Chow (n= 8), Cheese (n= 11).

**Figure 10.**
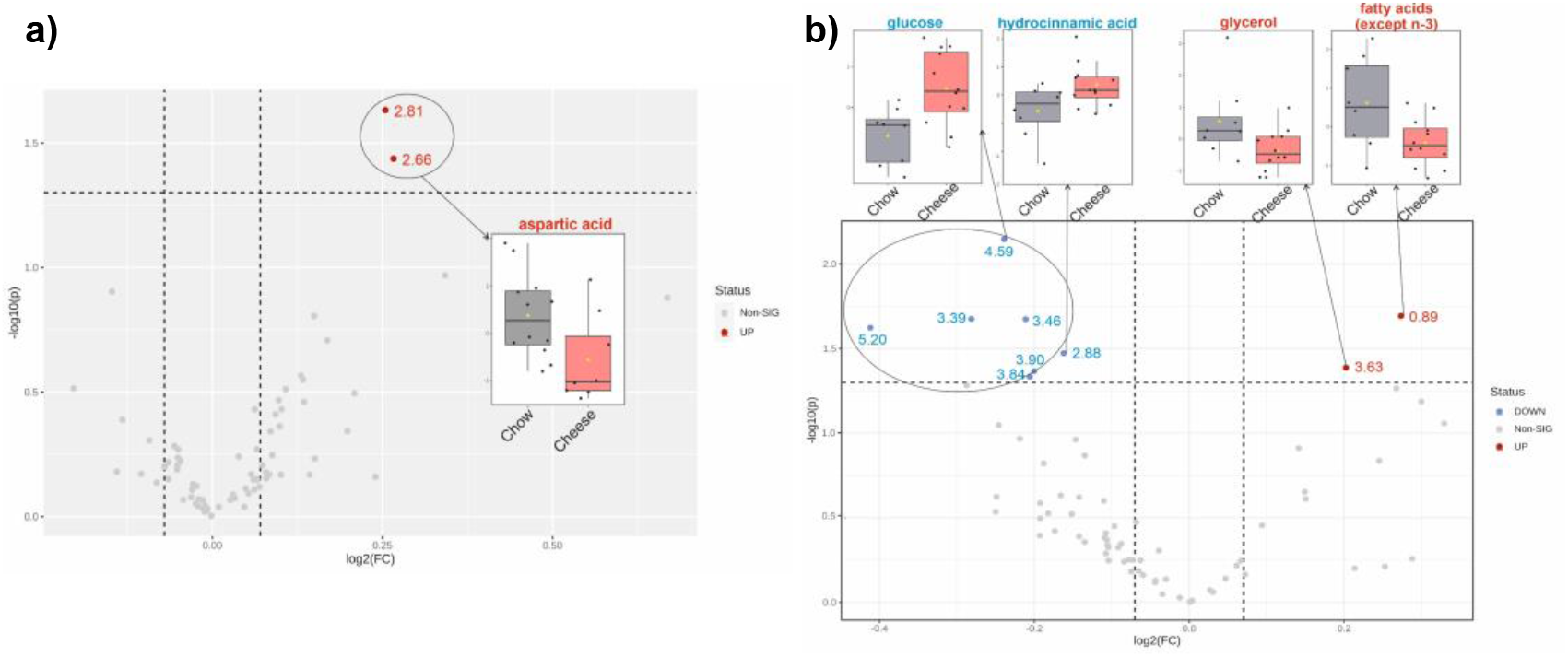
Volcano plots for (A) male and (B) female feces showing the differential metabolites between Chow and Cheese groups, displaying in ordinate the level of significant difference (− log10(p-value)) and in the abscissa the expression fold change (log2FC). All statistical valid FC of metabolites were considered according to Wilcoxon rank-sum tests (*p* < 0.05). Male volcano plot has n = 12 for Chow and n = 8 for Cheese; while female volcano plot has n = 8 for Chow, and n = 12 for Cheese.

Once we have detected the metabolites, a principal component analyses (PCA) was applied to detect any visual discrimination between groups. This PCA revealed thatonly in the female stool samples, a visual discrimination between Chow and Cheese groups was possible. Furthermore, supervised PLS-DA analysis showed the same result, a valid discrimination between Chow and Cheese groups but only in the female’s samples.

NMR peaks were analyzed deeply with univariate analysis, showing that there were some metabolites with significant fold-changes between Chow and Cheese groups for male and female groups. Those results are shown in the table 1. Furthermore, volcano diagram analyses showed an increase of aspartic acid and both FA (assigned to the CH_3_ terminal of all FA chains except those from omega-3 FA) in male stool samples, while glycerol in female feces for Chow group. In addition, glucose and hydrocinnamic acid were decreased in Chow group in female stool samples.

**Table 1.**
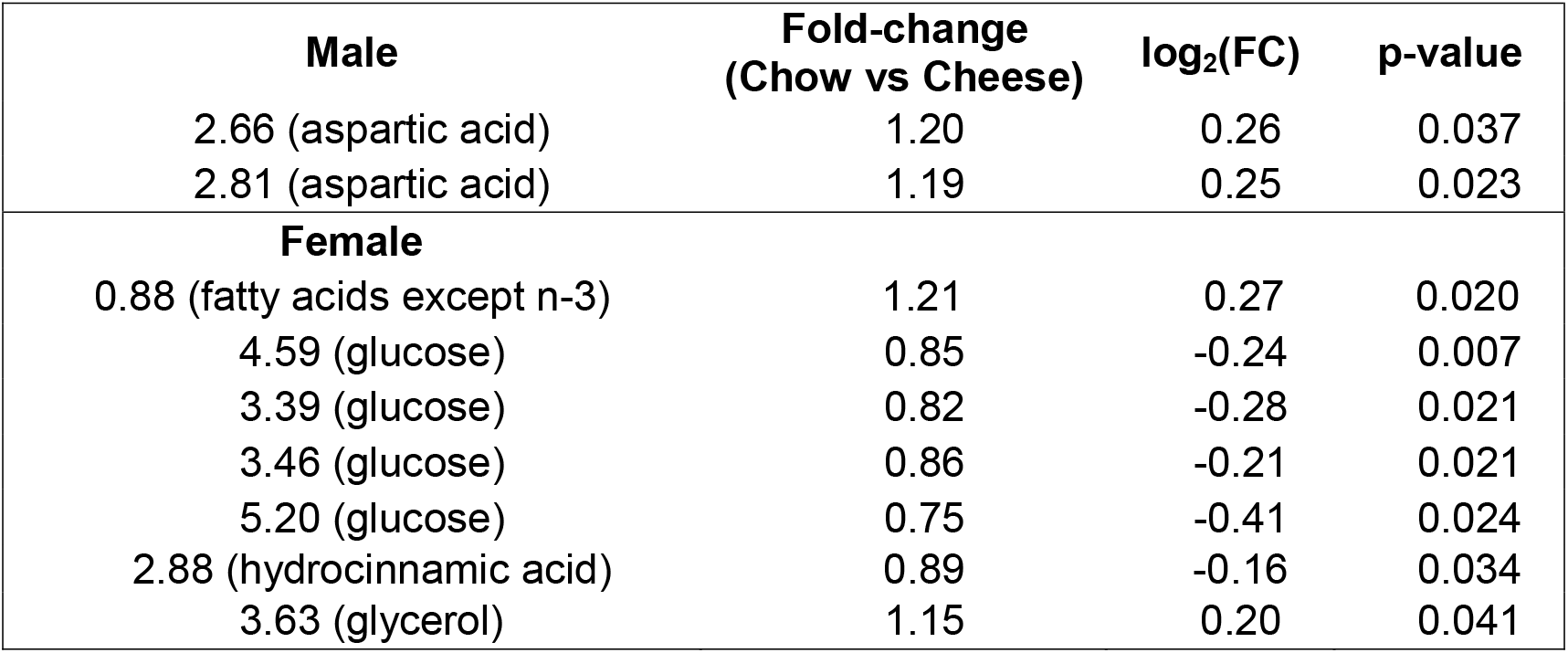
Fold-changes and *p*-values for differential NMR peaks between Chow and Cheese groups in male and female feces.

| Male | Fold-change<br>(Chow vs Cheese) | log<sub>2</sub>(FC) | p-value |
| --- | --- | --- | --- |
| 2.66 (aspartic acid) | 1.20 | 0.26 | 0.037 |
| 2.81 (aspartic acid) | 1.19 | 0.25 | 0.023 |
| Female |  |  |  |
| 0.88 (fatty acids except n-3) | 1.21 | 0.27 | 0.020 |
| 4.59 (glucose) | 0.85 | -0.24 | 0.007 |
| 3.39 (glucose) | 0.82 | -0.28 | 0.021 |
| 3.46 (glucose) | 0.86 | -0.21 | 0.021 |
| 5.20 (glucose) | 0.75 | -0.41 | 0.024 |
| 2.88 (hydrocinnamic acid) | 0.89 | -0.16 | 0.034 |
| 3.63 (glycerol) | 1.15 | 0.20 | 0.041 |

The NMR spectra showed that, in male stool samples, oleic acid was increased in the Chow group despite FA were not discriminating biomarkers (figure 11 and Table 2). This result may be due to the reduced fold-change obtained, that might be difficult to detect in NMR spectra where all FA chains overlap. In female samples, a 2-fold increase in lauric acid (a saturated FA) was present in Chow group. So, it is possible to conclude that this saturated FA was the responsible for the discriminatory difference in the integral of the CH_3_ terminal peak from all FA except *n-3* FA (that include saturated FA) previously found by NMR between Cheese and Chow groups.

**Figure 11.**
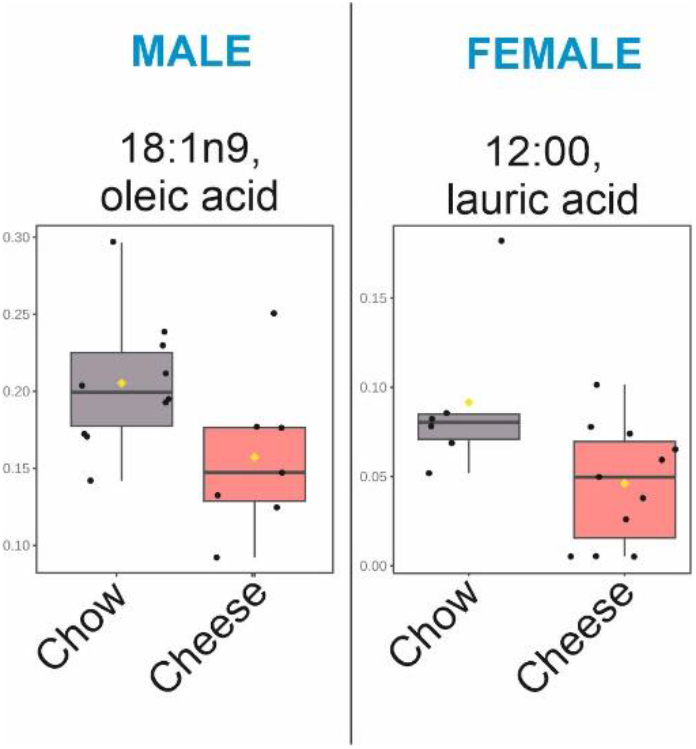
Box-plots, fold-changes and *p*-values for differential FA concentrations between Chow and Cheese groups in male and female feces by GC-FID. Oleic Acid n is as follows: Chow (n = 10), Cheese (n = 7); Lauric Acid n is: Chow (n = 6), Cheese (n = 11).

**Table 2.** Fold-changes and *p*-values for differential NMR peaks between Chow and Cheese groups in male and female feces for FA.

| **Male** | **Fold-change**<br>**(Chow vs Cheese)** | **log<sub>2</sub>(FC)** | **p-value** |
| --- | --- | --- | --- |
| 18:1n9 (oleic acid) | 1.3 | 0.38 | 0.045 |
| **Female** |  |  |  |
| 12:00 (lauric acid) | 2.0 | 0.99 | 0.032 |

## 4. Discussion

The present work showed increased premature behavior in the HFD group (Cheese) compared to the control. Interestingly, as can be seen in the rGT results, HFD is however able to modify impulsive choice, but it seems that this effect may be task-dependent. Regarding the neurochemical results, an increase and a reduction of BDNF relative expression was shown in males and females respectively. Furthermore, an increased relative expression of *DRD1* and *CB1* receptors can be observed. The ELISA’s results show that TNF-α and leptin levels change through development. The metabolomics analyses revealed that HFD exposure in adolescence is able to affect metabolism in adulthood. An additional summary of these results can be found in the Table S3.

Our results showed dramatic and long-term effects on impulsive behavior in rats exposed to HFD in terms of premature responses in tasks such as the 5-CSRTT and VDS. Remarkably, our results were unrelated to bodyweight, were no significant differences were found. These results are in accordance with previous studies like in Adams et al. (2015) where the diets (HFD, HSD) were administered during adulthood, while ours was administered during adolescence. Their results show that HFD performed more premature responses compared to the other groups. Literature also supports the idea that high motor impulsivity is related to higher high fat binge-like eating consumption (Anastasio et al., 2019), thus pointing to the possibility of the existence of a vulnerability in motor impulsivity after HFD consumption in adolescence. In addition, our findings are in accordance with Cussotto et al. (2018) review, where the authors show a relationship between diet-related factors and different behaviors.

In regard to choice impulsivity assessed with the delay responses in the DDT, no effect was found. Our results are in accordance with others like in Garman et al. (2021) where a HFD was provided *ad libitum* for 14 days and no differences in impulsive choice were found. In addition, Narayanaswami et al. (2014) studied the breakpoint of rats exposed at a HFD for 8 weeks; they divided the sample into obesity-prone (OP) and obesity-resistant (OR) according to body weight. No difference was found between the OP and OR groups. This long-term vulnerability seems to not affect all the impulsivity subtypes (Dalley, Everitt & Robbins, 2011) because effects were found in motor but not in choice impulsivity. Aside from the heterogeneous nature of the phenomenon, this difference can be explained by methodological variability across studies: in the study previously reported (Narayanaswami et al., 2014), they used data from the quartile 1 and quartile 4 (rats that gained the most weight vs rats that did not gain enough weight) while other studies did not use this approach. This division has been used as a suitable method for studying diet-induced obesity (DIO; Levin et al., 1997; Boustany et al., 2004), but, arguably, when the objective is not to directly study an obesity phenotype, subtle differences in inhibitory control deficit might be difficult to identify, or even fail to appear. Another explanation could be the differences regarding diet type (HFD, HSD, Caf; Jacques et al., 2019; Robertson & Rasmussen, 2017) and time that it is available (Moreira-Júnior et al., 2021; Veniaminova et al., 2020; Steele, Pirkle & Kirkpatrick, 2017). Steele et al. (2017) found that rats preferred the SS more than the LL in the DDT when they performed the task when off diet. However, their choice changed and they preferred the LL more than the SS when on HFD.

There is little data regarding clinical studies that analyze the relationship between obesity and impulsivity phenotype and, thus, the relationship between diet and impulsivity in humans. Some authors (Pan, Li, Feng & Hong, 2018) found a positive relationship between data of the body-mass-index (BMI) and cognitive instability and motivation impulsivity. On the one hand, when the phenomenon is assessed during adolescence, a positive relationship appears between people with higher fatty and sugary diets and sensation seeking and positive and negative urgency (Smith et al., 2021). On the other hand, when it is assessed after adolescence (youth), no differences were seen in impulsivity regarding BMI, but there existed an effect on motor impulsivity (Chamberlain, Derbyshire, Leppink & Grant, 2015). Finally, when this relationship is analyzed across adulthood the relationship is clearer in cognitive impulsivity than in motor impulsivity according to the meta-analysis performed by Amlung et al. (2016), but these differences can only be seen if there is an obese phenotype (Davis et al., 2011; VanderBroek-Stice et al., 2017).

Regarding compulsivity, our results show no deleterious effect of HFD. No differences were observed in perseverative responses on VDS, 5-CSRTT, and rGT. These results go in accordance with previous observations (Narayanaswami et al., 2014; Adams et al. 2015); however, some authors have reported differences in the Marble Burying Test (Moreira-Júnior et al., 2021; Souza-Gomes et al., 2018) developing DIO. These results were also seen in monkeys, where they made more perseverative responses in a reversal learning task when HFD was consumed *ad libitum* (Wilson & Howell, 2019). More research is needed to fully understand the relationship between compulsivity and diet. Kakoschke, Aarts and Verdejo-Garcia (2019) exposed that this relationship could be explained by: contingency-related cognitive flexibility, task/attentional Set-Shifting, attentional bias/disengagement, and the results of habit learning.

Intriguing long-term effects of a HFD consumption are seen in decision making assessed with the rGT in the present study. HFD rats tend to be more conservative in their responses. Seemingly, they tend to cope with less risk by making more premature responses in the most rewarded and less in the 50%-reward and 50%-punish options. This result is in discrepancy with the reported motor impulsivity tasks such as 5-CSRT task and VDS. Both tasks assess impulsivity/compulsivity with light-dependent response. The subject must respond to the light making a nosepoke where it has been shown for a specific period of time to receive a reward (Bari, Dalley and Robbins, 2008). However, even though the rGT is also a light-dependent task, it assesses impulsive decision making, not motor impulsivity. Some authors have considered the rGT as a waiting impulsivity task (Barkus et al., 2018; Tremblay et al., 2019); however, the task’s objectives can lead to a different consideration of its nature. In the 5-CSRTT, the animal only has to respond to a light in order to get a reward, while in the rGT, the animal must follow a specific decision making process in order to choose the option which fits the better (see methodology part). These differences might root on the rGT (Langdon et al., 2019) and 5-CSRT task (Bari, Dalley & Robbins, 2008) distinct methodologies. Moreover, we also found a difference in the percentage Choice-selection on P3 and in Choice-selection index showing that HFD were more conservative choices than control. Regarding clinical models, Navas et al. (2016) found that obese individuals made more riskier choices than controls, showing a possible link between risky decision making and obesity. These results may seem contrary, however, the influence of diets on decision making may be explained through two possible options. First, a reinforcement devaluation process could be present in our measure. Cycled-Caf diets showed reduced liking behavior to a sucrose solution at 2% (our test diet is 3% sucrose) (Martire, Westbrook & Morris, 2015). Second, the foraging strategies changed through the caloric value of the different diets. The influence of palatability on motivation to respond for non-caloric food and caloric food is different if the rats are food-deprived or not (Scheggi et al., 2013).

Regarding gene expression in RTqPCR, long-term alteration was seen in fold change in genes related to two of the main neurotransmitter systems that are related to food regulation. Long-term effects were detected in *DRD1* expression only in males, while no differences were seen in females. This result is in accordance with previous results in other brain areas like brainstem (Alsiö et al., 2014) and amygdala (Fülling et al., 2020), two components of the mesolimbic pathway (Koob & Volkow, 2010). The endocannabinoid system was also affected. Satta et al. (2018) found reduced levels of anandamide in frontal cortex and amygdala but higher levels in Nacb. On the other side, the 2-arachidonoyl glycerol was increased in HCC. This system has been proposed to be closely related with food rewards, and associated with the DA system (Martire et al., 2014). Moreover, differences in endocannabinoid tone were found in other mesolimbic pathways such as amygdala, caudate-putamen and hippocampus (Martire et al., 2014; Satta et al., 2018).

Some studies have found an effect of HFD on HCC morphology and function (del Olmo & Ruiz-Gayo, 2018; Valladolid-Acebes et al., 2013; Fernández-Felipe et al. 2021). To our knowledge, no studies have reported BDNF dimorphism in the same work as inhibitory control deficits. While BDNF expression increased in males, it decreases in females. It seems that females are more vulnerable than males in terms of long-term vulnerability after HFD consumption in adolescence. Some studies have shown that HFD and Caf is associated with reduced levels of BDNF in the HCC (Prochnik et al., 2022; Martire et al., 2014).

When assessing leptin concentrations, no differences were seen between groups. This effect is of significance since it seems to strengthen our findings of altered phenotype without differences in bodyweight, thus pointing to a vulnerability mechanism which is not related to body composition or obesogenic condition (Miller et al., 2014; DiLeone, 2009). The potential explanation of behavioral abnormalities being related to neuroinflammation cannot be confirmed either, since no differences were observed in TNF-α in serum or in RTqPCR for TNF-α. Future studies should be addressed to answer the question of which mechanism might underlie the present phenomenon, the role that neuroinflammation could play unrelated to TNF-α, and the contribution of other brain areas important to motor impulsivity, such as the nucleus accumbens.

On the one hand, previous research using NMR procedure detected deficits in gut metabolites in rodents with abnormal inhibitory control. Specifically, Abreu et al. (2022) found that compulsive-like rats, selected using the scheduled induced polydipsia (Moreno & Flores, 2012) had higher fatty acid levels (but not saturated) than the control group. In addition, glucose and glycerol levels were reduced in the compulsive-like group. Merchan et al. (2019) found a similar pattern: lower levels of glucose in the serum of the high compulsive-like group. Glucose influences decisions by indicating the body’s energy status, rather than merely acting as a source for replenishing cognitive processing efforts (Shechter & Schwartz, 2018). Wang & Huangfu (2017) tested this idea by giving some participants glucose at different dosages and asking them to complete an intertemporal delay discounting task. They results revealed a negative correlation between the rate of delay with blood glucose levels. This effect was only present in the glucose-feed group suggesting that the behavioral effects were related to hunger-reduction. Our results may go in accordance with their idea; a predictive mechanism of the glucose-insulin system for managing both metabolic and behavioral aspects of acquiring and distributing resources. Furthermore, Hui et al. (2014) showed that women with gestational diabetes (that have the glucose-insulin system unbalanced) have problems with adaptations to dietary management in a limited time period. Few information can be found about the relationship between FA and decision making. In a randomized control trial, Antypa et al. (2009) detected that after a supplementation with omega-3, the participants made fewer risk-aversive decisions tan the placebo group. Summarizing this information, glucose may affect decision making by modifying the strategies required to manage metabolic and behavioral outcomes.

On the other hand, the relationship between FA and inhibitory control deficits is still unknown. As Agostini et al. (2017) claimed in their review, the association between ADHD symptoms and n-3 FA (omega 3) represents a consistent finding among observational studies, but less evidence that link the other type of FA with ADHD symptoms can be found. Pase et al. (2015) showed that, after a HFD exposure, in an early development period, an hyperactive behavior could be seen after maturation. Our results complete those of Pase et al. (2015). We found that, after an adolescent HFD consumption, higher levels of FA were present in the exposed group, that revealed higher ADHD like behavior (increased motor impulsivity and risky decision making). However, most of the research found that feeding supplementation with omega 3 reduces ADHD-like behaviors in rodents and in humans (Agostini et al., 2017). Our results also may be related with this pattern, no effect of HFD was found on omega 3 FA, but the rest were affected (by sex or diet).

In summary, we found that HFD consumption in a critical developmental stage is related to long-term deficits in inhibitory control, specifically in impulsive-like behaviors. Moreover, this long-term vulnerability seems to differentially impact the different subcomponents of inhibitory control. In addition, HFD consumption affects PFC, thus dramatically interfering with mesolimbic pathway function. Furthermore, HFD exposure modifies the gut metabolic profile, affecting the FA, glucose, and other compounds related to different neurobehavioral outcomes. To our knowledge, this is the first study that has found long-term effects in impulsive behavior when HFD consumption is performed in adolescence. More research is needed to disentangle the specific mechanisms underlying these intriguing effects. To conclude, we have shown that exposure to a HFD in a developmental period (like adolescence) is able to create a long-term vulnerability in adulthood in all the domains analyzed. Moreover, it is worth noting the discriminatory effect that HFD has on impulsivity measures. It is able to impact some specific domains of impulsivity while leaving unchanged others. The relationship between HFD consumption and decision making needs more research due to their complexity.

## Acknowledgements

This work was supported by the following funding sources: UAL2020- CTSD2068 with FEDER I+D+I funds “Una manera de hacer Europa”; PND-2022l024 Plan Nacional sobre Drogas, Ministerio de Sanidad, Gobierno de España and PID2022-139286NB-I00 Proyectos de Generación, Gobierno de España and FEDER Funds.

## 5. Declaration of generative AI in scientific writing

During the preparation of this work the author(s) used ChatGPT3 in order to enhance readability and text understanding. After using this tool/service, the author(s) reviewed and edited the content as needed and take(s) full responsibility for the content of the publication.

