## Supplemental Material 1 for "“From Nutritional Patterns to Behavior: High-Fat Diet Influences on Inhibitory Control, Brain Gene Expression and Metabolomics in Rats”"

### **Supplementary 1 - Behavioral and biochemical procedures-**

**Behavior analysis**

### **1. *Variable Delay to Signal (VDS)***

VDS task procedure was based on Leite-Almeida et al. (2013). The task was conducted with the central hole of the 5-CSRT task panel. Animals were habituated to the test for two consecutive days. In the first session, the animals were placed in the operant chamber for 30 minutes with the chamber light and the food magazine light on, the remaining holes plugged with vinyl tape, and five dustless sweet pellets (pellets from now on; Noyes dustless reward pellets, TSE Systems, Germany) available in the magazine and the central hole. On the second session, animals were placed in the operant chambers with the lights on, the 5 holes unplugged, and pellets available in the magazine and the central hole. Ten consecutive training sessions (twice daily) followed the habituation phase, where the trial started with the house light on and one pellet. Once the trial started, a 3 second delay appeared with the house light on (delayed period), continued by a response-light of 60s (response period). Nosepokes made were either rewarded (if done during the response time; 60s; correct response) or punished (if done within the delay period; 3s; premature response) with a timeout (TO) period of five seconds in complete darkness. The inter-trial-interval (ITI) was set at 3 seconds. Each of the training sessions ended once the rat performed 100 trials or has been in the OC for 30 minutes. The test session was similar to the training phase, excepting the delay and the TO: a total of 120 trials (or ending after 60 minutes) were divided into different latencies in four blocks: 3 second (first and last 25) and randomly-occurring 6/12 second (70 middle) trials. Multiple premature nosepokes were allowed; these did not initiate the TO period. Along with premature responses, the following parameters were considered: total session time, total correct response, total omissions, total and mean response latency, total and mean reward latency for measuring task performance and perseverative responses at training and test conditions.

### **2. *Delay Discounting Task (DDT)***

For this task, two levers (A and B) were presented in the OC. If the rat pressed lever A, a small reward (smaller, sooner, SS; one pellet) would be given, while the rat pressed the B lever, a large reward (larger, later, LL; 4 pellets) would be given. The house and the food magazine light were used as a signal to start the trial, and a nose-poke was needed to initiate the lever presentation. When a response (press-response) was applied, the house light was turned off, the “clicker” sounded and the levers were retracted. The behavioral test was divided into six sets of three sessions each with different delays for receiving the reward. After two days of habituation, with the same reward for the small and the large, the phase of delay 0 started (set 1), receiving the large and small reward instantaneously. Session with delay 0 was followed by sessions with delay 5 (set 2), 10 (set 3), 20 (set 4) and 40 (set 5) seconds of delay. The last, set 6, was identical to the first one. The lever contingencies remained the same for every subject but were counterbalanced between rats. The trial ended with a response or the failure to make the nose-poke within 10 seconds. The variables chosen to analyze were the large reward choices and the lower reward choices. Each set consisted of 60 trials of which half were free choice (both levers available) and half were forced trials (where only one lever was available). Impulsivity was analyzed with the number of times that the rat chose the higher reward lever in the free choice trials.

### **3. *Five-Choice Serial Reaction Time Task (5-CSRTT)***

For this task, the complete panel of the 5-CSRT task was used. After a habituation phase where pellets were placed in the food magazine and in the 5 holes, rats were trained to respond to brief flashes of light presented in one of the five spatial locations randomly (Carli et al. 1983) within a daily session consisting of 100 discrete trials; task performance was stabilized after a mean of 12 sessions. Sessions started with house, food magazine light on and a pellet delivery. A nosepoke response triggered the trial start. After the trial started, the food magazine light was turned off and an ITI of 5 s started, and at its end a visual stimulus of 1-s duration (stimulus duration, SD) was presented randomly in one of the 5 holes. Responses to this hole within 5 seconds (known as the limited hold, LH) were considered as a correct response and rewarded with a pellet. The following response errors were considered: commission errors (response to the wrong holes), premature responses (those made before the light stimulus was presented); omissions (when not responding within the LH) and perseverative response (an additional response in a rewarded hole between a correct response and the pellet collection). If one of these errors were made (excepting perseverative response), a 5-s time-out period (house light off and not reinforcing) occurred. Each session consisted of 100 trials and 30 minutes; sessions ended if one of these conditions appeared. For improving task acquisition, the SD was reduced progressively from 8-s to 1-s. Thereby, in the first phase, SD = 8-s with LH = 10-s and ITI = 5-s; while in the following phases SD was reduced to 6, 2.5, 1.25 and 1-s with LH and ITI = 5-s. For advancing within the phases, the following criterion must be achieved: at least 50 correct trials with > 80% of accuracy and < 20% of omissions. The training was performed according to previous procedures with subtle modifications (Bari et al., 2008; Robbins, 2002). The following behavioral measures were recorded: accuracy (% correct trials = correct trials/correct trials + incorrect trials x 100), nº of incorrect responses, nº of omissions, nº of correct responses, nº of premature responses, nº of perseverative responses and latencies (to correct response; to incorrect responses and to rewards). Behavioral measurements were calculated by taking the mean of three consecutive sessions with stable performance (< 20% omissions; > 80% accuracy; SD = 1 s; ITI = 5 s).

### **4. *Rat Gambling Task (rGT)***

For this task, four of the five holes (the center hole was not used) of the 5CSRT task panel were used. After a habituation phase, where pellets were placed in the 4 holes and in the food magazine for 30 minutes, rats were first trained to perform a nose-poke response in one of the four places indicated by a 10 second light. If the response was done within this interval, the behavior was rewarded with one pellet. The location of the response-light varied among the holes chosen during all sessions, and session ended if the rat completed 100 trials or stayed in the operant chamber for 30 minutes. The training continued until the rats achieved the criteria of 50 correct responses with <20% of omissions and >80% of accuracy. The rats had to initiate the trial by a nose-poke response into the food magazine, which turned off the magazine light and started a 5 second delay. Any response performed within this delay period was considered as a premature response and punished with a 5 seconds TO (lights on). A lack of response within the 10 seconds was considered an omission. If the rat responded with a nose-poke to one of the response-light, reward was presented.

After the training phase, rats went under a forced-choice variant of the rGT, where only one response-light among the four in each trial was illuminated and the final contingencies were present: each hole was associated with a configuration of probabilities/magnitudes of reward (0.4/4 pellets, 0.5/3 pellets, 0.8/2 pellets, or 0.9/1 pellet) and punishment times (0.6/40 s, 0.5/30 s, 0.2/10 s, or 0.1/5 s), respectively. This ensured that the rats had the same experience with reward contingencies before the free-choice phase.

Finally, the animals went through a free choice phase where all holes were illuminated at the same time. The variables analyzed were choice preference, response latency and omissions across all the test sessions. For a detailed explanation of the paradigm, please see: Langdon et al (2019). The following variables were analyzed: percent premature responses; percent choice = (nº choices made for a pellet option)/(N responses made) x 100; choice score = (P1 + P2) - (P3 + P4); percent perseverative responses, percent correct and punished trials and latencies.

**Biochemical analysis**

**RTq-PCR**

Samples from prefrontal cortex and nucleus accumbens were used for gene expression analysis. Trizol method (Ambion) was used for RNA isolation. RNA total concentration was quantified by fluorometry with a Qubit (Invitrogen). Samples were normalized to 20 ng/uL, using this concentration for cDNA synthesis (20uL). These 20uL were diluted in 80uL of RNAse free water, and this dilution was used for the RT-qPCR. RT-qPCR was performed in microplates, which are composed of SYBR green master mix, primers, cDNA and nuclease-free water (total volume of 10uL). Within these microplates, samples were duplicated, run and analyzed in a thermocycler. Abnormal patterns were detected analyzing carefully Ct values and melting curves. Primers designed in exon sections of *gapdh* gene were used as housekeeping gene, while intron sections designed primers of *gapdh* were used for gDNA contamination assessment. The following genes were analyzed: TNF-α (forward: 5’-GGAGGGAGAACAGCAACTCC-3’; reverse: 5’-GCCAGTGTATGAGAGGGACG-3’; own design), GAD1 (forward: 5’-gtgagtgccttcagggagag-3’; reverse: 5’-cgtcttgcggacatagttga-3’; Perez-Fernandez et al. (2020)), BDNF (forward: 5’-ggtcacagcggcagataa-3’; reverse: 5’-ccgaacatacgattgggtag-3’; Perez-Fernandez et al. (2020)); TYRO (forward: 5’-CCTTCCAGTACAAGCACGGT-3’; reverse: 5’-TGGGTAGCATAGAGGCCCTT-3’; own design); DRD1 (forward: 5’-GCATGGCTTGGATTGCTACG-3’; reverse: 5’-CCAGTTGCTGCCTGGACTAA-3’; own design); DRD2 (forward: 5’-CAGTCGAGCTTTCAGAGCCA-3’; reverse: 5’-CCAATTCTCCGCCTGTTCACT-3’; own design). Samples with gDNA expression less than 30 (< Ct 30) and without abnormalities in the melting curves were used for further statistical analysis. Data transformation was performed as follows: mean Ct values of every sample were normalized to those obtained in the housekeeping gene (ΔCt) and, then, normalized to internal control (ΔCt), which is the average ΔCt of the Male-chow and Female-chow groups. To end, the ΔΔCt was transformed to obtain the fold change (2^ΔΔCt), which is the data picted. The present analysis method was already used by other researchers in our lab (Perez-Fernandez et al., 2022).

**ELISAs**

Blood samples were collected in all animals (chow and cheese) in eppendorfs without any coating. Samples were obtained from the lateral tail vein in both groups at two different times: PND75 and PND106. Previous blood extraction, rats were anesthetized with isoflurane in order to reduce stress of the tail-nick. Serum was separated by centrifugation (Sigma 3-18KS, Germany) of the blood samples at 3000 rpm (RCF = 800 xg) for 20 min at 4 ºC.; after separation, serum was stored at -20 ºC until analysis. Corticosterone levels were analyzed with an DetectX® enzyme immunoassay kit (K014-H1, Arbor Assays ™, Ann Arbor, USA). The inter- and intra-assay coefficients of variation were 6.56% and 6.65% respectively. Assay sensitivity was 6.71 pg/mL. Leptin levels were analyzed with an Abnova® enzyme immunoassay kit (KA0026-8, Abnova, Taipei, Taiwan). The inter- and intra-assay coefficients of variation were 0.006% and 5.25% respectively. Assay sensitivity was 50 pg/mL. TNF-α levels were assessed using a Sigma-Aldrich® enzyme immunoassay kit (RAB0479-1KT, Darmstadt, Germany). The inter- and intra-assay coefficients of variation were 0.003% and 6.14% respectively. Assay sensitivity was 25 pg/mL.

**Metabolomic analysis**

NMR spectroscopy was applied to analyze changes on the fecal metabolome on male and female rats between Chow and Cheese groups. Fecal analysis can offer valuable insights on the metabolic state of rats, their health sate, the impact of diet, and the involvement of the gut microbiota, which has a main role in various physiological processes and ultimately influence the composition and characteristics of feces. Thus, this non-invasive technique can help to elucidate the complex interplay between diet, metabolism, and gut health in rodent models.

Feces were frozen under -80 ºC until the moment of their analysis, and then were freeze-dried for 72 hours. For all samples, 20 mg of fecal matter were weighted and extracted with 0.7 mL of a 1:1 mixture of CH_3_OH-*d*_4_ and D_2_O KH_2_PO_4_ buffer 1.5M (pH 7.4) containing the sodium salt of 3-(trimethylsilyl)propionic-2,2,3,3-d4 acid (TSP, 0.1%, w/w) and sodium azide (NaN_3_, 90 *µ*M) as an enzyme inhibitor. Then, samples were submitted to sonication for 20 minutes, stirring for 10 minutes (600 rpm) and centrifugation for 5 minutes (13500 rpm). Finally, five hundred microliters of the supernatants were collected and transferred to 5 mm NMR tubes. A Bruker Avance III 600 MHz spectrometer operating at 600.13 MHz equipped with a thermostatted SampleJet autosampler of 500 positions and a 5 mm QCI quadruple resonance pulse field gradient cryoprobe was employed to conduct all ^1^H NMR experiments. ^1^H NMR spectra were obtained at 293 ± 0.1 K without rotation along with the suppression of the residual water signal employing a presaturation pulse sequence (Bruker 1D *noesygppr1d*). Acquisition parameters were set as following: 32 scans and 4 dummy scans, 65536 data points, spectral width of 20.5 ppm, acquisition time 2.73 s, relaxation delay of 5.0 s, FID resolution of 0.37 Hz and mixing time of 1000 ms. Lock was performed to the CH_3_OH-*d*_4_ frequency. All NMR spectra were bucketed by selecting and integrating all individual peaks observed in the region from δ_H_ 0.2 to 10.0 ppm using AMIX 3.9.15 (Bruker BioSpin GmbH, Rheinstetten, Germany). Normalization was performed on the data prior to statistical analyses through scaling the intensity of individual peaks to the total intensity obtained in the mentioned region. Regions that contain residual H_2_O suppression and methanol signal of δ_H_ 4.90−4.70 ppm and δ_H_ 3.31-3.34 ppm, respectively, were excluded from the bucketing and analysis. Metabolite assignments were performed thanks to information extracted through 2D NMR experiments (homo- and heteronuclear experiments) corresponding to ^1^H−^1^H COSY, ^1^H−^1^H TOCSY, ^1^H−^13^C HSQC and ^1^H−^13^C HMBC, along with the employment of some NMR databases (Chenomx and HMDB), and literature. The concentrations of different metabolites were calculated in relation to the inner standard (TSP) by means of the integration of peak areas from nonoverlapped signals.
