## Supplemental Material 2 for "“From Nutritional Patterns to Behavior: High-Fat Diet Influences on Inhibitory Control, Brain Gene Expression and Metabolomics in Rats”"

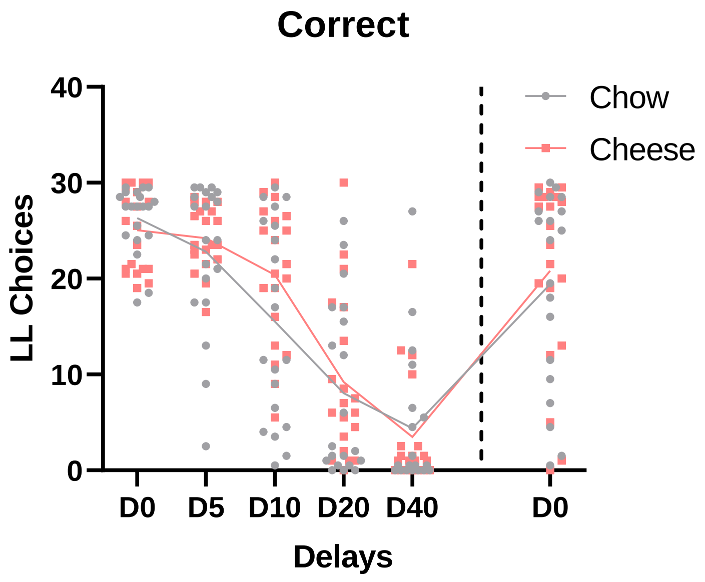


**Supplementary figure 1**. Graphical distribution of LL choices according to different delays. No diet effect was found comparing any of the delays used regarding the LL choices. Total n = 20 per group. Data is represented as mean ± SEM. *** $p < 0.001$

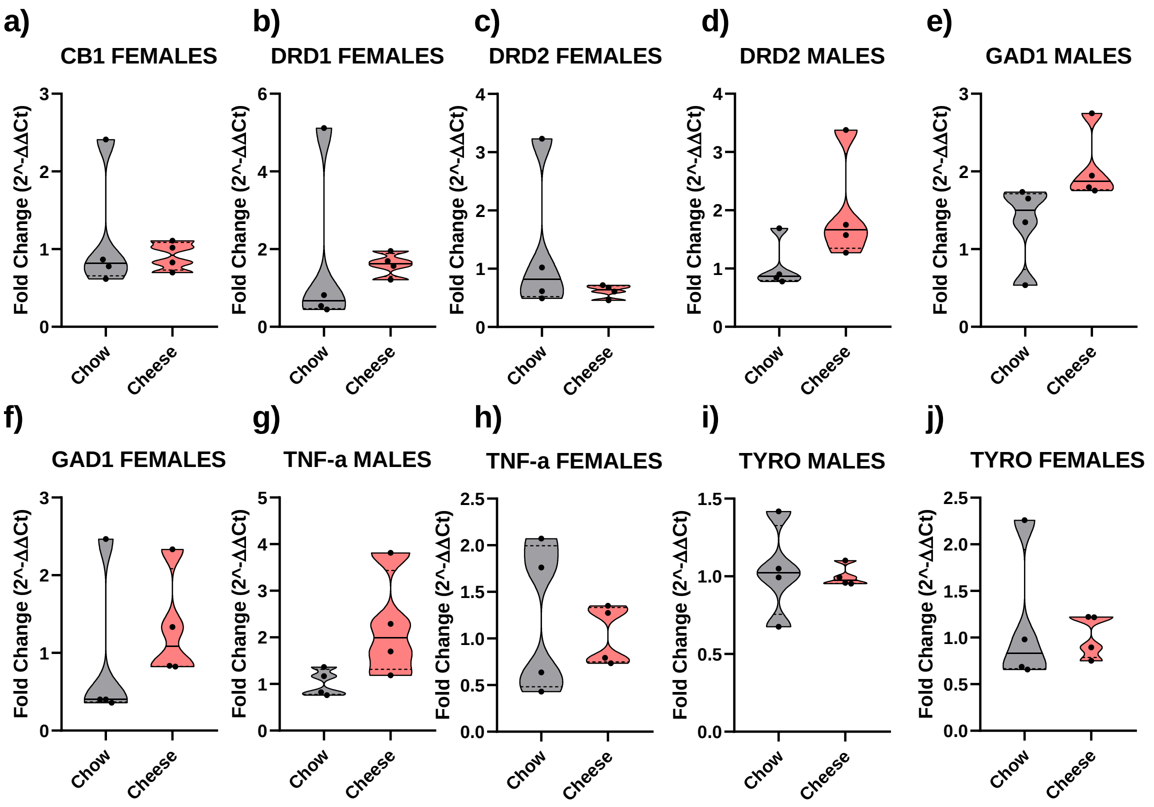


**Supplementary Figure 2.** Fold Change representation of the target genes analyzed with RT-qPCR in female/male rats (n = 4 in chow and n = 4 in cheese). Data is represented as mean ± SEM. * p < 0.05

**Table S1.** Metabolites identified in fecal extracts ($CD_3OD$:$D_2O$ $KH_2PO_4$ buffer 1.5 M at pH 7.4 (1:1, v/v) by $^1$H NMR. Homo and heteronuclear 2D NMR experiments were employed for the verification of the assignments, such as $^1$H-$^1$H TOCSY, $^1$H-$^1$H COSY $^1$H-$^{13}$C edited HSQC and $^1$H-$^{13}$C HMBC.

| **Metabolite** | **$\delta_H$ (ppm), *J* (Hz)** |
|---|---|
| Bile acids | 0.66 (s) |
| Fatty Acids | 0.88 (t, –C**H**$_3$, FA except *n*-3), 1.24 - 1.30 (m, –(C**H**$_2$)$_n$–), 1.55 (m, –C**H**$_2$–CH$_2$–COOR, all FA except DHA, ARA and EPA), 2.06 (m, –C**H**$_2$–CH=CH–, UFA), 2.34 (m,–C**H**$_2$–COOR, FA except DHA), 2.39 (m,–C**H**$_2$–COOR, DHA), 2.76 (m, =CH–C**H**$_2$–CH=, PUFA), 5.36-5.44 (m, –C**H**=C**H**–, UFA)* |
| Butyrate | 0.93 (t, *J*= 7.4 Hz), 1.58 (c, *J*= 7.4 Hz), 2.15 (t, *J*= 7.5 Hz) |
| Propionate | 1.08 (t, *J*= 7.6 Hz), 2.17 (c, *J*= 7.7 Hz) |
| Valine | 1.02 (d, *J* = 7.0 Hz), 1.06 (d, *J* = 7.0 Hz) |
| Isoleucine | 0.96 (t, *J* = 7.2 Hz), 1.03 (d, *J* = 7.0 Hz) |
| Leucine | 0.99 (d, *J*= 5.9 Hz), 0.98 (d, *J*= 5.9 Hz) |
| Threonine | 1.34 (d, *J*= 6.8 Hz), 3.54 (d), 4.24 (m) |
| Alanine | 1.49 (d, *J* = 7.2 Hz), 3.74 (c) |
| Acetate | 1.91 (s) |
| Glutamate | 2.05 (m), 2.13 (m), 2.39 (m), 3.72 (m) |
| Succinate | 2.41 (s) |
| Aspartate | 2.66 (dd, *J* = 9.3 Hz, 17.5 Hz), 2.81 (dd, *J* = 3.6 Hz, 17.5 Hz) |
| Hydrocinnamic acid | 2.45 (t, *J* = 8.5 Hz), 2.89 (t, *J* = 8.5 Hz), 7.28 (m) |
| Choline | 3.21 (s) |
| Betaine | 3.28 (s), 3.89 (s) |
| Glycerol | 3.55 (dd, *J*= 6.2 Hz, 11.5 Hz), 3.64 (dd, *J*= 4.6, 11.5 Hz) |

| | |
|---|---|
| Glycine | 3.50 (s) |
| Glucose | 4.60 (d, *J*= 7.5 Hz), 5.20 (d, *J*= 3.4 Hz) |
| Triacylglycerols (TAG) | 5.30 (b.s.) |
| Fumarate | 6.56 (s) |
| Tyrosine | 6.85 (d, *J* = 8.5 Hz), 7.19 (d, *J* = 8.5 Hz) |
| Phenylalanine | 7.34 (m), 7.41 (m) |
| Uracil | 5.75 (d, *J* = 7.7 Hz), 7.54 (d, *J* = 7.7 Hz) |
| Formate | 8.48 (s) |
| Nicotinate | 7.51, 8.29 (d), 8.59 (d), 8.99 (m) |

**Table S2.** Fatty acids identified by GC-FID and range of concentrations among all the sampels

| Chain | Name | Range of concentrations |
|---|---|---|
| ***Saturated*** | | |
| 6:00 | Caproic acid | 0 – 6.9% |
| 8:00 | Caprylic acid | 1.0 – 10.1 % |
| 10:00 | Capric acid | 1.1 – 12.9 % |
| 12:00 | Lauric acid | 0 – 11.4 % |
| 14:00 | Myristic acid | 1.8 – 10.0 % |
| 16:00 | Palmitic acid | 24.0 – 47.3 % |
| 18:00 | Stearic acid | 0 – 6.4 % |
| ***MUFA*** | | |
| 18:1n9 | Oleic acid | 7.3 – 27.0 % |
| 18:1n7 | Vaccenic acid | 0 – 14.0 % |
| 20:1n9 | Eicosenoic acid | 10.0 – 41.7 % |
| ***PUFA*** | | |
| 16:2n4 | All cis-9,12-hexadecatrienoic acid | 0 – 5.1 % |
| 16:3n4 | All cis-6,9,12-hexadecatrienoic acid | 0 – 6.1 % |
| 18:2n6 | Linoleic acid | 0 – 15.5 % |

**Supplementary table 3**

Table resume of the main findings feeding a high fat diet in a specific developmental period.

| | | | Males | | Females | |
|---|---|---|---|---|---|---|
| Assessments | | | Chow | Cheese | Chow | Cheese |
| Behaviour | Motor Impulsivity | VDS | ↓ | ↑ | ↓ | ↑ |
| | | 5-CSRTT | ↓ | ↑ | ↓ | ↑ |
| | Waiting Impulsivity | VDS | = | = | = | = |
| | | DDT | = | = | = | = |
| | Impulsive Choice | rGT | ↓ | ↑ | ↓ | ↑ |
| Biochemical variables | ELISAs | LEPT | ↑ | | ↓ | |
| | | TNF-α | ↓ | | ↑ | |
| | RT-qPCR | GAD1 | ↓ | ↑ (p = 0.084) | = | = |
| | | TYRO | = | = | = | = |
| | | TNF-α | ↓ | ↑ (p = 0.084) | = | = |
| | | DRD1 | ↓ | ↑ | = | = |
| | | DRD2 | = | = | = | = |
| | | CB1 | ↓ | ↑ | = | = |
| | | BDNF | ↓ | ↑ | ↑ | ↓ |
| | NMR/GC-FID | Saturated FA | ↓ | | ↑ | |

| | | | | | | | |
|---|---|---|---|---|---|---|---|
| | | MUFA | | = | | = | |
| | | PUFA | | ↓ | | ↑ | |
| | | Oleic acid | | ↑ | ↑ | = | |
| | | Lauric acid | | = | | ↑ | ↑ |
| | | NMR modeling | | No discriminative | | PLS-DA | |
| | | NMR peaks | Aspartic acid | ↑ | ↓ | ↑ | ↓ |
| | | | Fatty acids | ↓ | ↑ | ↓ | ↑ |
| | | | Glucose | ↓ | ↑ | ↓ | ↑ |
| | | | Hydrocinnamic acid | ↓ | ↑ | ↓ | ↑ |
| | | | Glycerol | ↑ | ↓ | ↑ | ↓ |
